# Systematic, high-throughput characterization of bacteriophage gene essentiality on diverse hosts

**DOI:** 10.1101/2024.10.10.617714

**Authors:** Jackie Chen, Erick D. Nilsen, Chutikarn Chitboonthavisuk, Charlie Y. Mo, Srivatsan Raman

## Abstract

Understanding core and conditional gene essentiality is crucial for decoding genotype-phenotype relationships in organisms. We present PhageMaP, a high-throughput method to create genome-scale phage knockout libraries for systematically assessing gene essentiality in bacteriophages. Using PhageMaP, we generate gene essentiality maps across hundreds of genes in the model phage T7 and the non-model phage Bas63, on diverse hosts. These maps provide fundamental insights into genome organization, gene function, and host-specific conditional essentiality. By applying PhageMaP to a collection of anti-phage defense systems, we uncover phage genes that either inhibit or activate eight defenses and offer novel mechanistic hypotheses. Furthermore, we engineer synthetic phages with enhanced infectivity by modular transfer of a PhageMaP-discovered defense inhibitor from Bas63 to T7. PhageMaP is generalizable, as it leverages homologous recombination, a universal cellular process, for locus-specific barcoding. This versatile tool advances bacteriophage functional genomics and accelerates rational phage design for therapy.

## Introduction

Bacteriophages (or phages), Earth’s most abundant biological entity, represent unparalleled genetic diversity. Yet, despite their ubiquity, much of this diversity remains functionally uncharacterized—even in the most well-studied phages^1–5^. While functional genomics in phages has lagged, functional genomics in microbes has made great progress. In microbial systems, genome-wide gene essentiality screens have been pivotal in functionally characterizing genes and uncovering molecular processes of life. For example, gene knockout screens in model microbes such as *Saccharomyces cerevisiae* and *Escherichia coli* revealed key genetic factors that affect cellular processes such as DNA repair, antimicrobial resistance, metabolism, and growth^6–13^.

These genome-wide knockout screens have identified broadly essential genes critical for viability and uncovered conditionally essential genes—those essential only in specific contexts. Conditionally essential genes often provide insight into how organisms adapt to various environmental challenges by tuning their physiological responses^14–17^. Phages must balance this need for environmental adaptability with limited genome size, making the characterization of conditionally essential genes particularly important^18^. To uncover conditionally essential genes in phages, gene essentiality screens must be performed across diverse host environments. Such an approach would mirror the ecological diversity that phages naturally encounter, providing insights into their evolutionary strategies and adaptability.

However, translating the successful approaches in model microbes to phages has proven challenging. Factors such as the host-dependent life cycle of phages, the absence of general selectable markers, and the lack of effective genome manipulation tools have impeded the creation of comprehensive conditional gene essentiality maps, even for well-characterized phages^19,20^. This knowledge gap restricts our understanding of phage biology, including their host interactions, life cycles, and applications in biotechnology and phage therapy^21–24^.

Current methods for investigating phage gene essentiality, including classical gene knockouts^25,26^, chemical mutagenesis^27^, iterative recombination-driven genome reduction^28^, and CRISPRi^29,30^, have significantly advanced our understanding of phage biology. Yet, these approaches have critical limitations. Classical genetic techniques can become impractical given the sheer number of genes in phage genomes (up to 100s)^2,3^. Chemical mutagenesis is untargeted, making it difficult to pinpoint causal mutations. Iterative genome reduction and CRISPRi are more systematic but come with their own challenges—iterative reduction requires laborious serial passaging, and CRISPRi suffers from polar effects which limits its ability to resolve gene essentiality in operons, a common feature in phages^29,31^. Moreover, iterative reduction and CRISPRi cannot screen on intractable hosts, which represent untapped reservoirs for revealing conditionally essential genes.

To overcome these limitations, we introduce **PhageMaP (<u>P</u>hage <u>h</u>igh-throughput <u>a</u>pproach for <u>g</u>ene <u>e</u>ssentiality <u>M</u>apping <u>a</u>nd <u>P</u>rofiling)**, a platform that generates pooled, barcoded phage knockouts for high-throughput screening of conditional essentiality. PhageMaP leverages Cas9 and recombinase to systematically interrogate genetic elements through homologous recombination (HR). Using pooled single-guide RNAs (sgRNAs) and barcoded donor sequences, PhageMaP synthesizes genome-wide barcoded knockouts in a single culture that can be tested across diverse environmental contexts, such as different host strains or growth conditions. Unlike methods based on transcriptional or translational repression, PhageMaP operates at the DNA level, avoiding polar effects and providing higher resolution in characterizing gene essentiality.

We apply PhageMaP to generate genome-wide barcoded knockout libraries for model and non-model coliphages, T7 and Bas63 (JohannRWettstein)^32^. We test these libraries against a panel of clinically relevant and common laboratory *E. coli* strains, as well as 33 *E. coli* strains harboring unique anti-phage defense systems^33^. PhageMaP successfully recapitulates long-established findings, including the essentiality of core structural and replication proteins. Using PhageMaP, we generate gene essentiality maps across hundreds of genes in the model phage T7 and the non-model phage Bas63, on diverse hosts. These maps uncover broad patterns of gene essentiality and provided fundamental insights into genome organization, gene function, and host-specific conditional essentiality. Further, we uncover phage genes that either inhibit or activate eight anti-phage defenses leading to new mechanistic hypotheses. We transfer one newly discovered anti-phage inhibitor from Bas63 to T7 to improve T7 infectivity against the *ppl* defense system, demonstrating how PhageMaP datasets can be leveraged for rational design of synthetic phages. Moreover, by assessing the libraries in various conditions, we assign new functions to both uncharacterized and characterized genes, broadening our understanding of phage biology. Together, our results highlight the diverse adaptations used by phages to thrive in different host environments – insights which can only be attained through the investigation of conditional essentiality. PhageMaP should be generalizable to other phages across different host backgrounds because it employs a universal process, homologous recombination, and a versatile gene editing tool, Cas9, for library creation^34^.

### PhageMaP creates barcoded genome-wide knockout libraries in phages

PhageMaP is carried out in three steps: 1) barcoded donor library synthesis using oligonucleotide pools, 2) donor cassette integration using *in vivo* Cas9-RecA-mediated recombination (Fig. 1A), and 3) deep sequencing to obtain quantitative gene essentiality mapping across conditions (Fig. 1B). We considered both *in vitro* and *in vivo* methods for creating barcoded libraries. *In vitro* methods have diminished generalizability because of bottlenecks from transformation efficiency or genome size^35,36^. In contrast, an *in vivo* HR approach leverages a conserved, universal cellular process^34^. We used Cas9-mediated HR to achieve site specific barcode insertion during phage infection^37^. We reasoned that this *in vivo* approach would be widely applicable to many phage-host systems, and as a proof of principle, applied PhageMaP to the well-studied coliphage T7.

**Figure 1.**
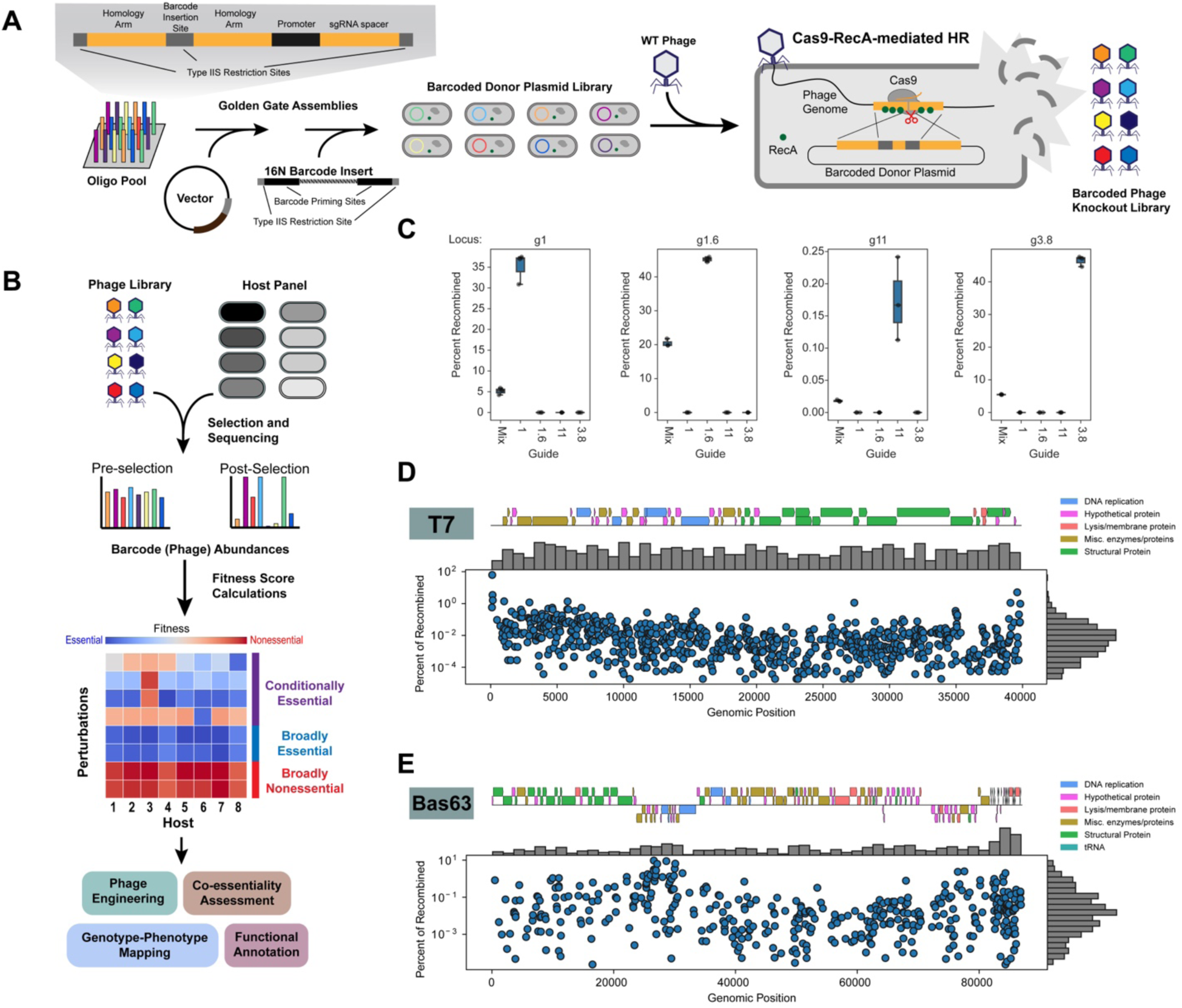
PhageMaP creates barcoded genome-wide knockout libraries in phages. **(A)** Scheme for generating barcoded phage knockout libraries. Oligos containing target-specific 95nt homology arms and sgRNAs are cloned into a high-copy number vector. 16N barcodes are subsequently added to generate a barcoded donor plasmid library. This library is infected by WT T7 and Cas9-RecA-mediated homologous recombination (HR) inserts barcodes into designated loci. **(B)** Screening of the PhageMaP libraries on diverse hosts. The pre- and post-selection abundances of phage variants are quantified with deep sequencing. Each variant is then scored to create context-specific gene essentiality profiles. **(C)** Validation of barcode insertions into four T7 genomic loci. Cas9-RecA-mediated HR was carried out using 1 of 4 sgRNAs or a mix of the 4 sgRNAs in pooled format. The four genomic loci were sequenced, and the percentage of recombined phages was determined by finding the ratio of insertions to total reads. **(D)** Distribution and mapping of the T7 PhageMaP library. **(E)** Distribution and mapping of the Bas63 PhageMaP library.

The donor library is used as the repair template for Cas9-RecA-mediated HR to generate the barcoded phage knockout library (Fig. 1A). RecA was added to boost recombination efficiency (Supplementary Fig. 1)^38^. Each donor containing a sgRNA spacer and site-specific homology arms is cloned into a vector where a random barcode is subsequently inserted. Barcodes are mapped based on the adjacent homology arms using short-read sequencing, thus completing the genotype-phenotype linkage. Upon recombination, the barcode is inserted into the target locus to disrupt the residing gene.

After recombination, the resulting barcoded phage knockout library can then be challenged on a panel of susceptible hosts. Pre- and post-selection barcode counts are used to calculate fitness scores and assess conditional essentiality across tested hosts (Fig. 1B). Mutants in essential regions should show lower fitness post-selection and have lower scores, while those in nonessential regions persist and have higher scores Because the same pre-selection phage library is used for each condition, biases that may arise from using different starting libraries of variants are accounted for.

To validate that PhageMaP can generate insertions at targeted loci, we assessed the recombination efficiencies for four loci in T7 individually and in pooled format. We tested two essential loci, *g1* (T7 RNA polymerase) and *g11* (tail tubular protein), and two reportedly nonessential loci encoding hypothetical proteins, *g1.6* and *g3.8*^27,28^. When targeted individually, the greatest recombination frequency occurred at a locus when the on-target sgRNA is used (0.1- 48%), and no recombinants were detected with the other off-target sgRNAs (Fig. 1C). Phage knockouts of essential genes are possible if the gene product is synthesized before Cas9 restriction. For instance, we observed ∼35% recombination of *g1* even though T7 RNA polymerase is essential (Fig. 1C). In pooled format, high fidelity recombination occurred at targeted loci albeit at a lower per-gene frequency (0.02-22%) (Fig. 1C). Thus, with our donor plasmid design and Cas9-RecA-mediated HR, we can efficiently and precisely insert barcodes into targeted loci of both essential and nonessential genes in parallel.

We next generated genome-wide libraries in T7 (Fig. 1D) which has a 40kb genome with about 60 genes^39^. We designed donor sequences for every 25-nucleotide window with an available sgRNA, resulting in a theoretical library size of 1084 sgRNAs encompassing both genic and intergenic regions. After assembly, the barcoded plasmid library contained 941 sgRNAs (85.8% theoretical) (Supplementary Fig. 2A). Variants were lost likely due to low initial abundance or toxicity. The barcoded phage library dropped to 779 sgRNAs (71.9% theoretical) after recombination (Supplementary Fig. 2A). After sequencing the barcoded phage library, we recovered 7447 unique barcodes (∼9-10 barcodes per sgRNA on average) and found that all 60 genes were represented in the library (Supplementary Figs. 2B-C). Using ddPCR, we estimated the percentage of recombinants in the population to be ∼2% (Supplementary Fig. 2D). We observed significant skew in the phage library toward the termini of the phage genome likely due to the repeats in this region promoting recombination (Fig. 1D)^40^.

To demonstrate generalizability, we also created PhageMaP libraries in a non-model phage, Bas63 (Fig. 1E). Bas63 is an 87kb phage encoding 161 genes^32^. To create a more uniform distribution of sgRNAs per gene, we selected 2-4 sgRNAs for each gene to create a theoretical plasmid library with 627 sgRNAs. After construction of the donor plasmid library, 473 sgRNAs were present (75.4%) (Supplementary Fig. 2A). Upon recombination, we obtained 8337 unique barcodes (∼20 per sgRNA on average) for 418 sgRNAs (66.7%) and recovered 149 out of 161 genes (Supplementary Figs. 2A-C). The fraction of recombinants was drastically higher in Bas63 (∼30%) than for T7 (∼2%). A possible explanation for this discrepancy is the absence of a RecBCD inhibitor in Bas63 which can impair the creation of single-stranded DNA for DNA repair through HR (Supplementary Fig. 2D)^41^. Compared to the T7 library, the Bas63 library is significantly less biased towards specific variants since no phage variant made up more than 10% of the total mapped barcodes (Fig. 1E).

In summary, PhageMaP comprehensively and precisely generates disruptions across the phage genome in regions that are either essential or nonessential and genic or intergenic for both model (T7) and non-model (Bas63) phages.

### PhageMaP recapitulates established biology and uncovers novel insights in a model phage

We selected T7 from the *Autographiviridae* family because its well-documented biology will help validate the results of our screen. Additionally, using T7 will challenge PhageMaP to uncover novel insights for a thoroughly researched phage. We screened the barcoded phage library by infecting target hosts and quantitatively scoring variants post-infection (See Methods). Here we mainly describe gene essentiality in relative terms (i.e. more or less essential), but binary labels can be applied by setting thresholds. Mutations in genes that improve phage fitness but are not critical for viability will appear more essential than mutations in benign regions. During analysis, overlapping genic regions are treated as unique “pseudogenes” with separate fitness scores. In T7, ten pseudogenes were found. Finally, the intergenic loci were treated as individual genes.

Though T7 is well-studied, comprehensive conditional gene essentiality maps for T7 do not exist. We applied the phage library on a panel of eleven *E. coli* strains – seven are common laboratory strains (BL21, BL21DE3, BW25113, MG1655, DH10B, S17, and LE392), two strains are clinical isolates from urinary tract infections (UTI33 and UTI46)^42^, and two are antibiotic-resistant strains from the ECOR panel (ECOR4 and ECOR13)^43^. Across the eleven hosts, we obtained 682 fitness measurements for T7 genes (Fig. 2A). Fitness scores were highly correlated among replicates (Pearson’s r: 0.90-0.98), indicating the barcode readout is robust and reproducible (Supplementary Fig. 4A). We performed unsupervised clustering of the normalized scores across all strains to group genes based on their gene essentiality profiles (Fig. 2A). Highly nonessential (dark red) and essential (dark blue) regions each comprise about 15% of total T7 genes, while the other genes are distributed throughout a continuous essentiality scale (Fig. 2A). The two large, equally sized clades distinguishing essential (blue) and nonessential (red) genes suggest half of the genes in T7 are essential (Fig. 2A). Overlaying the essentiality profiles with the T7 genome map revealed co-localization of genes with similar essentiality (Fig. 2B). For instance, essential genes *g1* to *g2.5* are grouped, while nonessential early genes up to *g0.7* are grouped (Fig. 2B). Evolutionarily, this grouping is likely preserved to maintain synteny of T7 genes with coordinated or dependent functions^3^.

**Figure 2.**
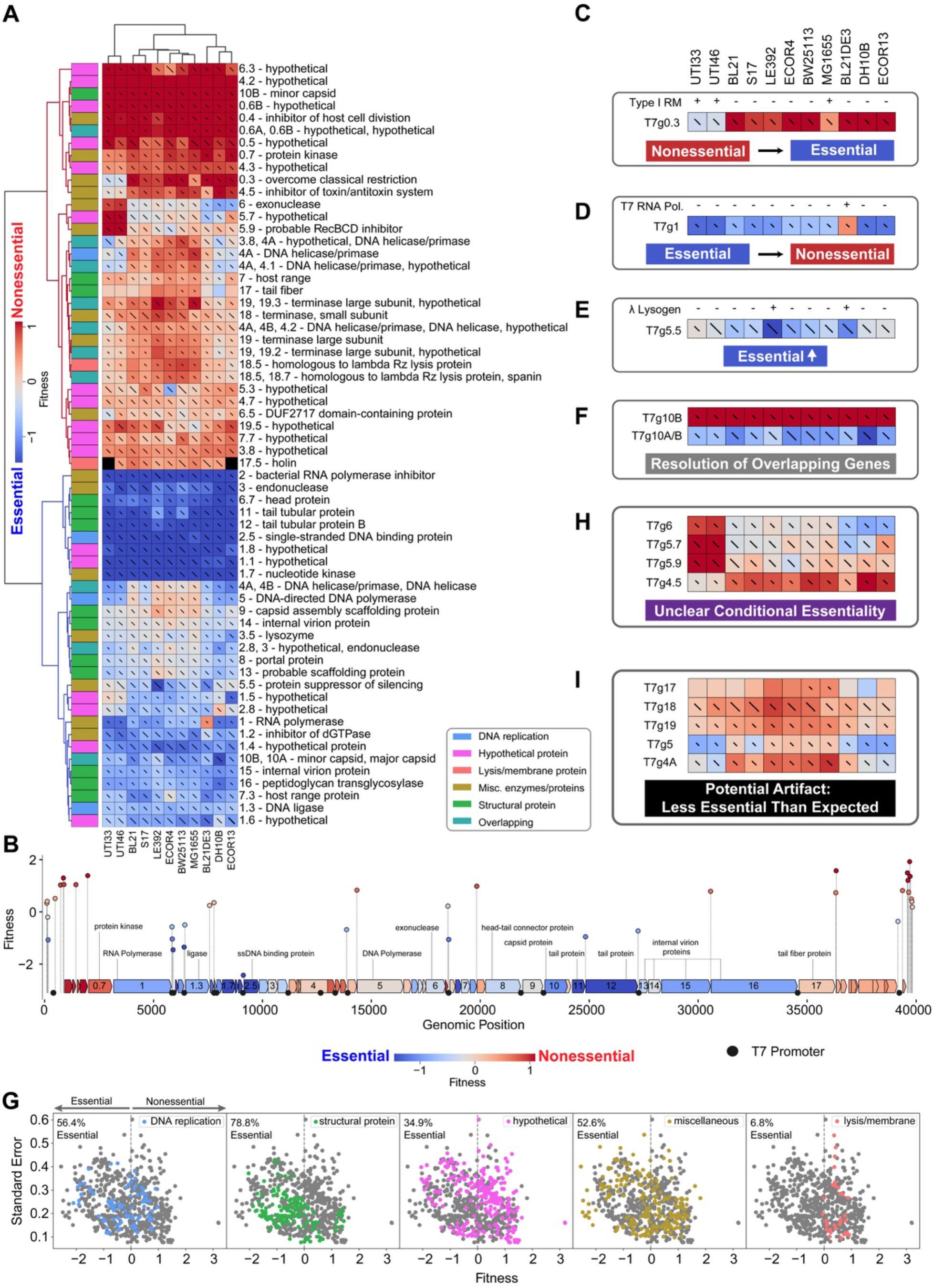
PhageMaP recapitulates established biology and uncovers novel insights in model T7 phage. **(A)** Heatmap of the T7 fitness scores after Z-normalization within each of the 11 conditions. To better emphasize differences outside the two extremes of the data, a 10^th^ percentile floor and 90^th^ percentile ceiling were imposed for the color scaling. The lines through each cell represent the standard error (SE) where a full line represents SE≥1. Clustering was performed using the Euclidean metric and Ward linkage method. Metadata describes general classification of each gene. **(B)** Plot mapping the genomic coordinates and scores for intergenic perturbations. The points (intergenic variants) and blocks (genes) are colored using the same scale as (A). The gene scores used for coloring are the medians of scores from each tested condition. Instances from (A) of **(C)** generally nonessential genes that become more essential. The presence of intact Type I restriction-modification systems was determined by used PADLOC^121^. Instances from (A) of **(D)** generally essential genes that become more nonessential, **(E)** moderately essential gene becoming more essential, and **(F)** resolution of overlapping genetic elements. **(G)** Scatter plots showing the fitness scores and standard errors for each fitness measurement for the 11 hosts. Points corresponding to each classification are highlighted by color. Dotted lines represent the threshold for “nonessential” and “essential” genes. Instances from (A) of **(H)** unclear conditional essentiality and **(I)** potential artifacts involving characterized genes.

We found that the PhageMaP results are consistent with prior knowledge of T7 biology and controls (Supplementary Fig. 4B). Genes involved in host takeover, nucleotide metabolism, and DNA replication (1.1-1.8, 3, 3.5, 4A/B, 5, 5.5, and 6) and structural proteins (6.7, 7.3, 9, 10A/B,11, 12, 13, 14, 15, 16) are generally clustered together in clades with high essentiality (Fig. 2A)^27,28^.

Highly dispensable early genes (0.3, 0.4, 0.5, 0.6A/B, and 0.7), hypothetical genes (4.2, 4.3, and 4.7), and gene 4.5 (inhibitor of toxin-antitoxin system) are clustered in groups with low essentiality. Gene 0.3, encoding the Ocr protein which protects phage DNA from restriction-modification (RM) systems^44^, is more essential for strains UTI33, UTI46, and MG1655 where the Type I RM system is intact (Fig. 2C)^45^. In addition, gene 1 (T7 RNA polymerase) is less essential in BL21DE3, a strain that contains endogenous T7 RNA Polymerase which can complement the loss of this crucial enzyme in T7 (Fig. 2D). Next, gene 5.5, which enables growth on lambda lysogens^46^, is more essential in LE392 and BL21DE3, the two strains containing a lambda prophage (Fig. 2E). The targeted approach of PhageMaP helps resolve overlapping genes. For instance, *g10A* (major capsid protein) fully overlaps with *g10B* (minor capsid protein), which has a 53-amino-acid extension beyond *g10A*’s stop codon. Loss of *g10B* reportedly doesn’t affect T7 viability provided *g10A* is present^47^. The higher fitness scores for *g10B* perturbations but the overlapping *g10A/B* regions across all tested strains support this (Fig. 2F). These results demonstrate that PhageMaP produces biologically meaningful gene essentiality patterns.

We further examined global patterns using a compilation of fitness profiles from the 11 strains by first classifying all T7 genes into 5 broad categories (DNA replication, structural, hypothetical, miscellaneous, and lysis/membrane) and then quantifying the percentage of scores where genes within a category are essential (fitness score < 0) (Fig. 2G). Most scores for structural (78.8%) and DNA replication genes (56.4%) were found to be essential. Hypothetical and miscellaneous proteins had a wide range of fitness scores (Fig. 2F). Many of these proteins serve generally essential roles for robust phage growth (e.g. nucleotide metabolism and host take-over)^48–50^ while others can serve conditionally essential roles in specific contexts (e.g. evasion of host defense)^41,44,51^. Lysis and membrane-related genes were mostly nonessential (6.8% essential). Although surprising as many of these proteins have roles in lysis and release of phage progeny, previous reports support the dispensability of those genes^52^. Our findings highlight the differences in T7 gene essentiality even within similar categories.

PhageMaP’s ability to directly edit the DNA of the phage enables interrogation of intergenic regions in ways CRISPRi screens, which rely on transcription or translation, cannot^29,30^. We obtained 364 fitness measurements for intergenic regions in T7 across the 11 strains. Seven intergenic variants located within DNA synthesis and class III gene clusters did not appear in the pre-selection phage library, presumably due to their roles in regulating the expression of essential genes. Regions near or at T7 promoters driving expression of essential genes (e.g. *g2.5*) are essential (Fig. 2B). Each of the three *E. coli* promoters which drive expression of the early genes is nonessential individually, possibly due to functional redundancy (Fig. 2B). Most loci at the genome ends of T7 also appear to be nonessential. The two exceptions are positions 142 and 162, near the end of the left terminal repeat (Fig. 2B). Because the direct terminal repeats play a role in DNA replication, concatenation, and packaging, these positions may contain critical elements^53^. In summary, the essentiality of intergenic genetic elements is generally consistent given their genomic location.

Some genes have discordant scores across the tested strains, indicative of conditional essentiality. For example, genes 6 (exonuclease), 5.7 (hypothetical protein), and 5.9 (RecBCD inhibitor) are less essential in UTI strains, which contain two intact prophages^54^ that may compensate for their absence (Fig. 2H, Supplementary Fig. 4C). Conversely, gene 4.5 is more essential in the UTI strains which express the CcdAB toxin/antitoxin (TA) system. Because gp4.5 inhibits the antiviral sanaAT TA system through lon protease inhibition^55^, we speculate that gp4.5 may inhibit the CcdAB TA system whose activity is also controlled by lon-dependent proteolysis^56^. These variations in essentiality suggest that further investigation could reveal previously unknown interactions.

Some unexpected fitness measurements may be the result of artifacts from the pooled planktonic nature of the screen (Fig. 2I). For example, *g17* encodes the tail fiber, a crucial structural protein for receptor recognition, but appears more nonessential than other tail proteins (e.g. genes 11 and 12). Previous studies also observed the nonessentiality of the T7 tail fiber in planktonic cultures due to complementation by free tail fiber proteins expressed from intact phages^27,57^. We posit that *g18* (terminase small subunit) and *g19* (terminase large subunit), encoding important enzymes for packaging the phage DNA into empty proheads, appear nonessential due to similar complementation. Genes 5 (DNA polymerase) and 4A (helicase/primase) are critical for phage DNA replication^58^ but appear to be nonessential. To determine if the persistence of these variants is due to free-floating DNA in the lysate, we DNase I-treated the lysate to remove signal from non-encapsulated phage DNA but found no difference (Supplementary Fig. 4D). Complementation by endogenous host enzymes, such as the *E. coli* DNA Polymerase I which is structurally similar to the T7 DNA Polymerase^59^, is possible, but these interactions have not been described in prior studies.

Taken together and despite the skew in the initial library, PhageMaP creates comprehensive gene essentiality maps of T7, encompassing both genic and intergenic regions across different hosts, to provide a global overview of conditional gene essentiality.

### PhageMaP creates gene essentiality profiles for a non-model phage

To demonstrate generalizability and scalability of our approach, we applied PhageMaP to characterize gene essentiality in Bas63, a recently isolated natural Myovirus phage belonging to the *Felixounavirus* subfamily^32^. Bas63 has more than double the genome size and almost triple the gene content of T7. Investigating phages with larger genomes may provide a wealth of insights into phage biology. Across 4 susceptible *E. coli* hosts (BW25113, DH10B, MG1655, and S17), we obtained 593 fitness measurements for 149 genes (Fig. 3A). The replicates from all four strains have a strong correlation (Pearson’s r: 0.80-0.89) (Supplementary Fig. 5). Unsupervised clustering of the genes based on their scores resulted in a clear formation of a clade belonging to putative essential genes consisting of structural and DNA replication genes and separate clades belonging to nonessential genes categorized as a mix of hypothetical, miscellaneous, and tRNA genes (Fig. 3A). Clustering revealed that about two-thirds of Bas63 genes are nonessential (Fig. 3A). While this might indicate Bas63 is burdened with deadweight, the observation of conditional essentiality within nonessential genes—such as hypothetical genes gp114 and gp116—suggests that these genes may constitute a “toolbox” for thriving in varied environments.

**Figure 3.**
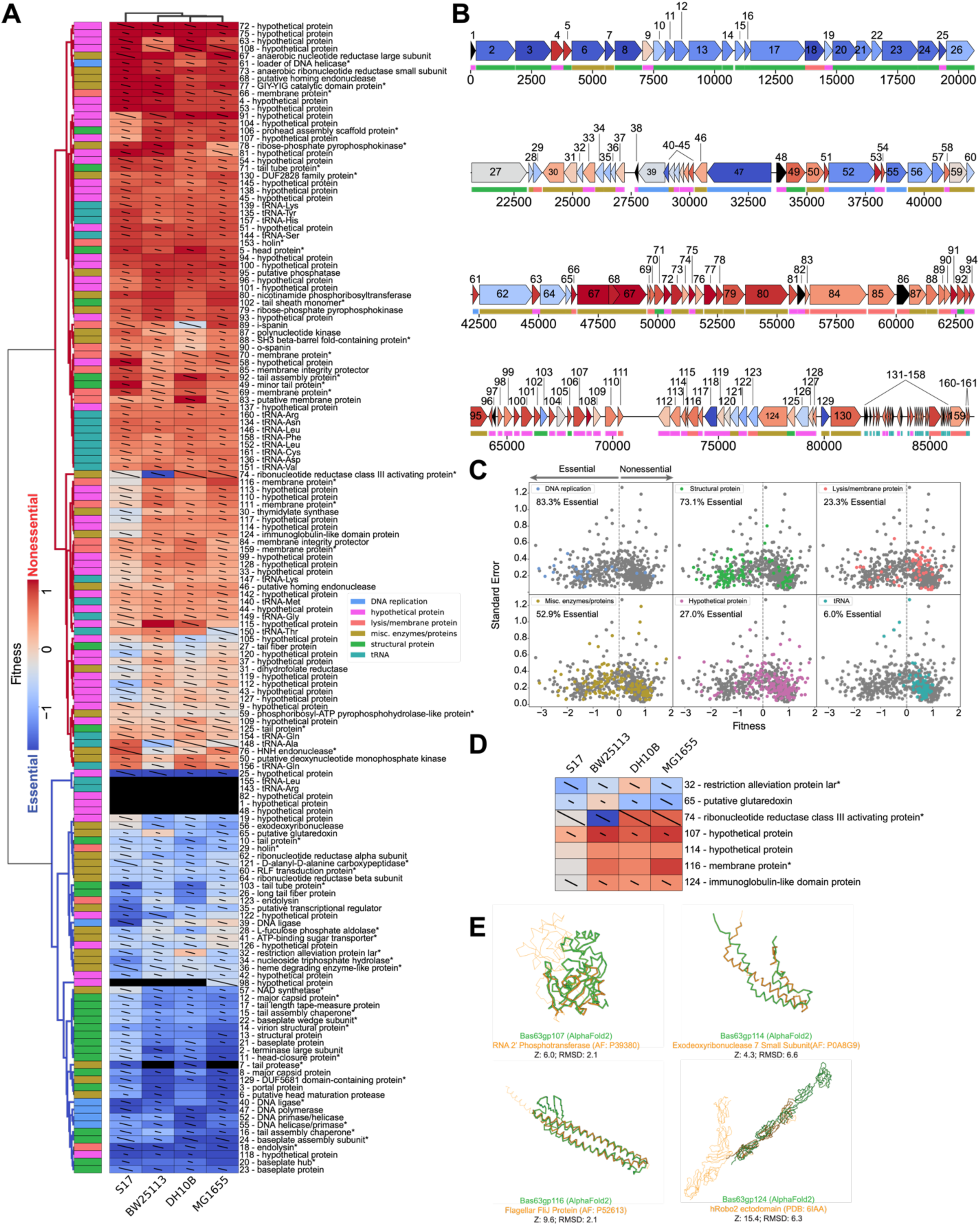
PhageMaP creates gene essentiality profiles for a non-model Bas63 phage. **(A)** Heatmap of the Bas63 fitness scores after Z-normalization within each of the 4 conditions. To better emphasize differences outside the two extremes of the data, a 10^th^ percentile floor and a 90^th^ percentile ceiling were imposed for the color scaling. The lines through each cell represent the standard error (SE) where a full line represents SE≥1. Clustering was performed using the Euclidean metric and Ward linkage method. Metadata describes general classification of each gene. Black boxes indicate no data was obtained for the gene in the corresponding condition. A black dot at the top right corner of a cell indicates only one replicate was obtained. Asterisks at the end of gene names indicate genes which were manually annotated after BLASTp search. **(B)** Bas63 genome architecture colored by the median fitness score for each gene from the tested conditions. Color scaling is the same as (A). **(C)** Scatter plots showing the fitness scores and standard errors for each fitness measurement for the 11 hosts. Points corresponding to each classification are highlighted by color. Dotted lines represent the threshold for “nonessential” and “essential” genes. **(D)** Patterns of conditional essentiality from (A). **(E)** Structural similarities between conditionally essential proteins and proteins found within the AlphaFold database and the PDB.

A prevailing theme in phage genomes is genetic mosaicism where loci accumulate from horizontal gene transfer events over time^2,3^. Like T7, we hypothesized that clusters of genes migrate as individual units based on similar function. Since function dictates gene essentiality, we investigated if mosaicism is also evident from the gene essentiality profiles in Bas63 by overlaying the median fitness score for each gene over the genome map. We observed segments that are uniformly essential (e.g. positions 0 to ∼21,000) or nonessential (e.g. positions ∼46,000 to ∼66,000) (Fig. 3B). Genes in the essential segments primarily code for structural proteins such as the tail- and capsid-related proteins. These genes likely mobilize as a cohesive unit, because their products often require interactions with other structural proteins^3^. The nonessential segments contain genes encoding hypothetical proteins, lysis proteins such as holins, and enzymes such as ribonucleotide reductases and pyrophosphokinases. Genes for these proteins may function dependently with neighboring genes hence their co-localization. The diversity within these segments suggests that phages allocate highly flexible regions of the genome for acquiring additional functions. These flexible regions may be separate from the essential regions to prevent disruption of core genes. Outside these segments, we observed mosaicism in regions containing interspersed small hypothetical genes with varying essentiality that may function more independently (e.g. positions ∼66,000 to ∼80,000) (Fig. 3B).

We next binned Bas63 genes into 6 broad categories (DNA replication, structural, miscellaneous, hypothetical, lysis/membrane, and tRNA) and found gene essentiality varies within these categories. Across the 4 strains, DNA replication and structural genes were scored as essential (fitness < 0) 83.3% and 73.1% of the time, respectively (Fig. 3C). Miscellaneous enzymes and hypothetical proteins exhibited a mix of essentiality. These proteins play roles in various cellular processes including nucleotide and nicotinamide metabolism and redox reactions. Lysis/membrane proteins were mostly nonessential. Unlike T7, Bas63 encodes tRNAs, which were mostly nonessential (Fig. 3C). Because phage tRNAs are thought to compensate for host tRNA deficiencies or provide resistance to viral defense systems comprising of tRNA nucleases^60,61^, Bas63 tRNAs are likely components within the conditionally essential toolbox that is leveraged in specific contexts.

In order to identify instances of conditional essentiality, we analyzed genes with discordant scores among the four tested strains (Fig. 3D). Gene 32, with 90% homology to the restriction alleviation protein Lar, is less essential in DH10B than the other three strains. Lar has canonical anti-restriction activity against Type I RM systems^62^, but these systems are not intact in S17-1 and BW25113. However, S17-1, BW25113, and MG1655 contain Type IV restriction systems absent in DH10B, suggesting that gene 32 may help evade these defenses. Gene 65 (putative glutaredoxin) and gene 74 (ribonucleotide reductase activator) have distinct essentiality in BW25113, indicating possible differences in nucleotide metabolism regulation in this strain.

S17-1 has a distinct gene essentiality profile clustered separately from the other strains (Fig. 3A). Among other genetic differences, S17-1 is the only tested strain that contains transfer (*tra*) genes involved in formation of the pili and conjugation^63^. We postulated mechanisms for four genes (*g107*, *g114*, *g116*, and *g124*) where scores for S17-1 were divergent and more essential than in the other strains (Fig. 3D). Gp107 shares slight structural homology with the NAD-binding C-terminal domain of RNA 2’ phosphotransferase^64^, suggesting Gp107 may facilitate NAD- dependent processes that improve Bas63 infectivity in S17-1 (Fig. 3E). Gp114 roughly superimposes on the small subunit of *E. coli* exodeoxyribonuclease 7, which in excess, can counter cellular toxicity of the large subunit^65^ (Fig. 3E). We speculate that large subunit toxicity may be a mechanism for phage defense in S17-1 and is overcome by expression of gp114 during phage infection. Gp116 shares sequence and structural homology with phage and bacterial membrane proteins including the flagellar FliJ protein (Fig. 3E). During lysis, gp116 may be necessary to efficiently disrupt the unique membrane architecture from pili formation^66^. Gp124 contains immunoglobulin (Ig) domains that can be aligned to the Ig domains of Robo2, which interacts with cell surface structures^67,68^ (Fig. 3E). Because *tra* genes can alter the membrane surface^69^ and Ig-like domains are thought to facilitate phage adsorption through cell surface interactions^70^, gp124 may have a role in Bas63 adherence to S17-1. Although further experiments are necessary to confirm these speculations, we show that these hypotheses can be made in part due to PhageMaP results.

In summary, we demonstrated that PhageMaP can be applied to a non-model *E. coli* phage. We showed that most Bas63 genes are dispensable in our tested conditions, but these genes likely serve beneficial functions when challenged to the appropriate condition.

### PhageMaP identifies anti-phage defense counters and triggers in a non-model phage

Beyond a gene essentiality tool, PhageMaP can be extended to investigate mechanistic relationships in specific biological systems. In recent years, there has been an explosion of newly discovered and enzymatically diverse anti-phage defense systems^33,71–74^. To counter bacterial immunity, phages have mechanisms to evade or block antiviral effector functions^73,75,76^. However, the specific interactions between phage gene products and most anti-phage defense systems are poorly understood. PhageMaP provides an opportunity to systematically and efficiently interrogate all genes within a phage genome to find novel interactions with anti-phage defense systems. Phage gene products can be “counters” or “triggers”. “Counters” inhibit host anti-phage defense to improve phage fitness and are more essential under anti-phage defense. Triggers activate host anti-phage defense to reduce phage fitness and are more nonessential under anti-phage defense. To uncover the relationship between phage genes and anti-phage defense, we challenged the Bas63 PhageMaP library to an *E. coli* panel of 33 anti-phage defense systems discovered and constructed by Gao *et al*^33^ (Supplementary Fig. 6A). In contrast to the previous sections where bacterial strains have many genetic differences that make it difficult to pinpoint strain-specific characteristics responsible for a phenotype, this anti-phage defense panel provides a controlled background where each defense system is expressed under the same genetic context, allowing for the definitive identification of causal interactions.

We challenged the Bas63 PhageMaP library on a 13-member subset of the 33-member defense panel that was susceptible to wildtype Bas63 because phage titers after infection were high enough to generate sufficient product for sequencing. We obtained 1911 fitness measurements for 12 strains, with reasonable correlation (Pearson’s r >0.58) between replicate pairs for all strains except RADAR2 which was removed from further analysis due to low correlation (r=0.2 in one replicate pair) (Supplementary Fig. 6B). To identify genes most likely to play a role during anti-phage defense, we statistically compared the Z-normalized scores of each gene in the presence of the defense system and with the no defense control. A total of 286 statistically significant fitness scores (p < 0.05, minimum difference=0.5) were identified across 12 defense strains and 56 genes (Fig. 4A). After adjusting for multiple testing (q < 0.1), 39 fitness scores remain across 9 strains and 11 genes. While each defense system has a unique essentiality profile, certain genes became either more essential (e.g. *g79* and *g80*) or more nonessential (e.g. *g40* and *g123*) in multiple defense systems, which indicates phage counters and triggers are not exclusive to specific defense systems (Fig. 4A). The varying degrees of essentiality within genes suggests counters and triggers have a range of effectiveness dependent on the defense system (Fig. 4A).

**Figure 4.**
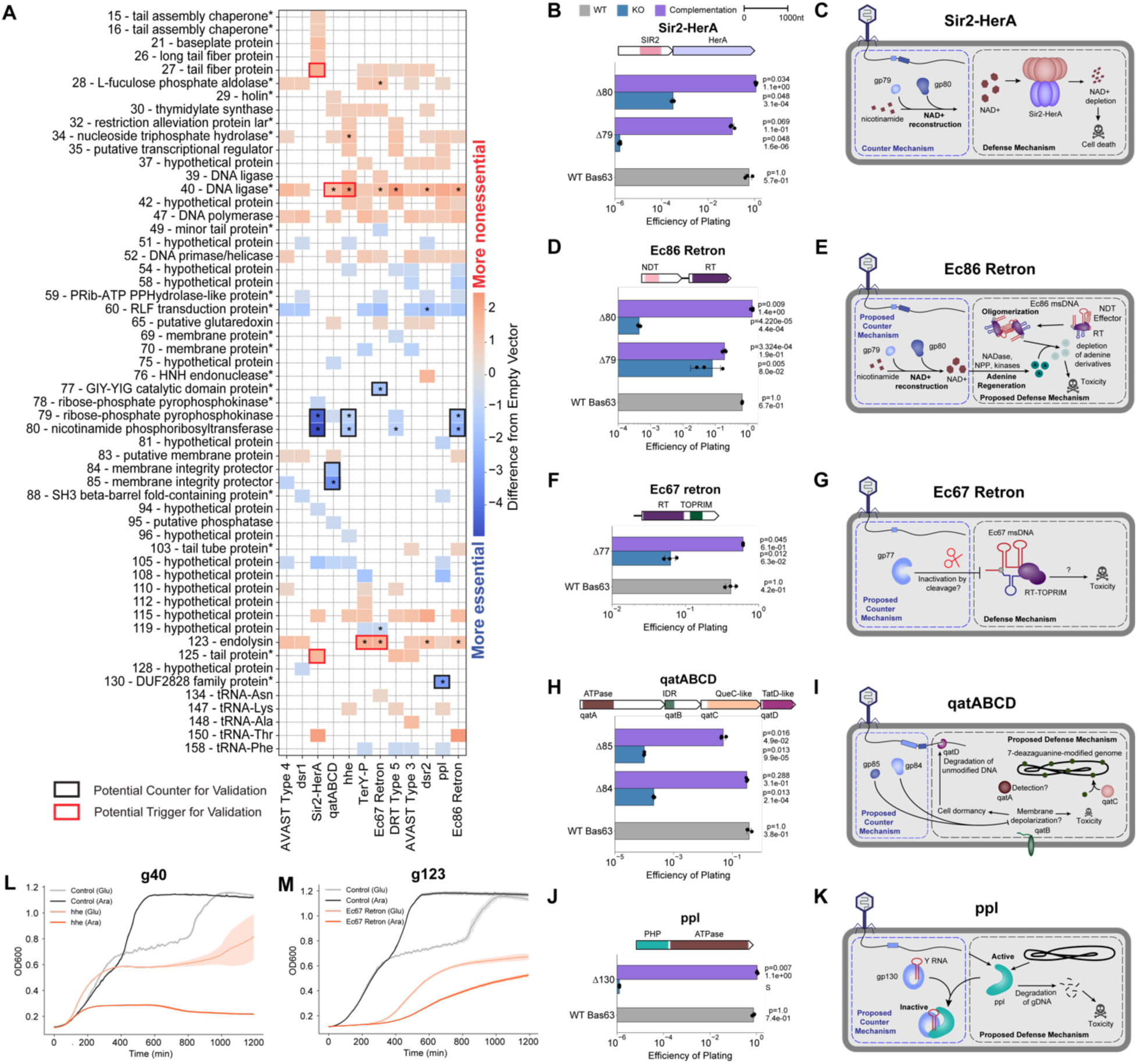
PhageMaP identifies anti-phage defense counters and triggers in Bas63 phage. **(A)** Plot showing all identified uncorrected significant hits (Welch’s t-test, p<0.05) and significant hits after multiple testing correction with the Benjamini-Hochberg method (q<0.1, denoted by an asterisk in each tile) with a score that is at least a difference of 0.5 from the control. The color of the tiles represents the difference in Z-normalized fitness scores between with defense and the control. Black boxes indicate targets deem to be potential counters to a defense system and subjected for secondary validation. Red boxes indicate targets deem to be potential triggers for a defense system and subjected for secondary validation. Asterisks at the end of gene names indicate genes which were manually annotated after BLASTp search. **(B)** Efficiency of plating (EOP) measurements for Bas63 knockouts with the Sir2-HerA host. Average titers on the control strain were first determined (3 technical replicates). Each point on the plot represents a technical replicate for EOP using those calculated averages as the denominator for each titer measurement on the defense host. Bars represent the mean of three replicates, and error bars denote the standard deviation. Hypothesis testing was performed pairwise with each knockout and WT. P-values (Welch’s t-test) and means are annotated above the bars. Complementation host contained the defense system and a plasmid expressing deleted gene. **(C)** Mechanisms of defense and counter-defense for Sir2-HerA. **(D)** EOPs for Ec86 retron calculated the same way as (B). **(E)** Proposed mechanisms of defense and counter-defense for Ec86 retron. **(F)** EOPs for Ec67 retron calculated the same way as (B). **(G)** Proposed mechanisms of defense and counter-defense for Ec67 retron. **(H)** EOPs for *qatABCD* calculated the same way as (B). **(I)** Proposed mechanisms of defense and counter-defense for *qatABCD*. **(J)** EOPs for *ppl* calculated the same way as (B). Since plaques were too small to count, an ‘S’ was annotated in lieu of a plaquing efficiency. The plotted efficiency is the highest Bas63Δ130 dilution for which zones of clearing were observable and clear on a spot plate. **(K)** Proposed mechanisms of defense and counter-defense for *ppl*. **(L)** Growth curves for expression of *g40* in cells containing no exogenous defense or the *hhe* system. 1 x 10^7^ cells were either repressed with 0.2% glucose (uninduced) or induced with 0.1% L-arabinose and grown for ∼20hrs. Central line represents the average of 3 replicates and the shaded flanking regions represent the standard deviation. **(M)** Growth curves for expression of g123 in cells containing no exogenous defense or the Ec67 retron system. Experiment was performed and plot was constructed the same way as (H).

We set out to clonally validate 10 counters and 6 triggers. For Sir2-HerA, *g79* and *g80* are potential counters (Fig. 4A). To validate this, we deleted each gene individually and assessed the ability of the mutants to infect Sir2-HerA cells compared to wildtype phage. Deletion of either *g79* or *g80* led to a 10⁴-10⁶ reduction in plating efficiency in Sir2-HerA cells, which was restored when the genes were complemented (Fig. 4B). Sir2-HerA is a bipartite phage defense system that induces abortive infection through depletion of NAD+, a vital cofactor^77^. Genes 79 and 80 both encode enzymes involved in NAD+ biosynthesis^78^. This suggests that Bas63 contains NAD+ synthesizing enzymes that regenerate NAD+ upon depletion by Sir2-HerA (Fig. 4C). Recently, a separate study validated that *g79* and *g80* are responsible for reconstituting NAD+ through novel pathways^79^.

Genes 79 and 80 are counters to the Ec86 retron and *hhe* defense systems, albeit with lower effectiveness than in Sir2-HerA (Fig. 4A). Ec86 retron plaquing efficiency was reduced ∼10-fold and ∼10^3^-fold when *g79* and *g80* were removed, respectively. These deficiencies are at least partially recovered upon complementation (Fig. 4D). Ec86 retron consists of a non-coding RNA (ncRNA), a nucleoside deoxyribosyltransferase-like (NDT) effector, and a reverse transcriptase (RT) that synthesizes a RNA-DNA hybrid molecule, multicopy single-stranded DNA (msDNA)^80,81^. These components assemble into oligomers to deplete adenine derivatives and inhibit phage replication^81^. NAD+ synthesized by gp79 and gp80 can be consumed to regenerate depleted adenine^82,83^ (Fig. 4E). For the *hhe* defense system, which consists of a single gene containing helicase and nuclease domains, deletion of either counter yielded a reduction in plaque size instead of plating efficiency (Supplementary Fig. 7A-B). We performed pooled planktonic competition assays with WT Bas63 and Bas63Δ*g79* and found that Bas63Δ*g79* is less fit than wildtype Bas63 in *hhe* cells, but this effect was negated after complementation (Supplementary Fig. 7C). This suggests that the *hhe* defense system may also involve NAD+ or adenine depletion. Taken together, these data highlight the convergence of phage counterstrategies to distinct anti-phage defense systems and the ability of PhageMaP to identify co-essential genes that function within a shared pathway.

Gp77 is a counter to Ec67 retron (Fig. 4A). Deletion of gp77 resulted in a 10-fold reduction in plating efficiency, which was restored when gp77 was complemented on a plasmid (Fig. 4F). The Ec67 retron consists of a reverse transcriptase-endonuclease fusion gene and a ncRNA that produces msDNA^33,84^. Gene 77 contains a GIY-YIG catalytic domain which is involved in DNA repair, transfer of retroelements, or degradation of foreign DNA^85^. Previously, it has been shown that degradation of the msDNA precursor is a mechanism for retron inactivation^86^. We propose that gp77 cleaves and inactivates the Ec67 msDNA to evade anti-phage activity (Fig. 4G).

For *qatABCD*, PhageMaP identified *g85* and *g84* to be counters (Fig. 4A). Deletion of either of these two genes resulted in a 10^4^-10^5^ reduction of plating efficiency that is largely restored after gene complementation (Fig. 4H). QatABCD comprises of four genes: QatA, which contains an ATPase domain; QatB, an uncharacterized protein with structural similarity to transmembrane proteins (Supplementary Fig. 7D); QatC, which contains a QueC domain possibly involved in DNA modification^87^; and QatD, a TatD nuclease which has a role in DNA fragmentation^88^. The exact mechanism for *qatABCD* defense is unknown. Genes 85 and 84 encode membrane integrity protector proteins that share high sequence homology (>80%) to RIIA/B lysis inhibition proteins found in coliphage T4. RIIA and RIIB inhibit RexAB-mediated membrane depolarization to allow T4 infection of lambda lysogens^89^. We propose that *qatABCD* inhibits Bas63 replication through a similar mechanism where membrane depolarization by QatB induces cell death or dormancy and activates QatD degradation of unmodified phage genomes (Fig 4I). The host genome is protected due to DNA modification by QueC (Fig 4I). QatA may be responsible for regulation of the anti-phage defense process, as is common with ATPases in other systems^90–92^.

PhageMaP identified *g130* as a *ppl* counter. Deletion of gene 130 severely reduced Bas63 plaque size, but this phenotype was reversed upon gene complementation (Fig. 4J). Ppl is an uncharacterized anti-phage system consisting of a single gene containing an ATPase domain and a Polymerase and Histidinol Phosphatase (PHP) domain with putative phosphodiester- hydrolyzing function^93^. Gp130 shares high structural similarity to Ro60, a protein that associates with noncoding Y RNAs to regulate RNA processing and decay through enzymes such as exoribonucleases (Supplementary Fig. 7E)^94^. Since Y RNAs structurally resemble tRNAs^94^ and *g130* is found directly upstream of a stretch of genes annotated as tRNAs in the Bas63 genome, some tRNA genes in Bas63 are possibly Y RNAs that can associate with gp130 for regulatory functions. We propose that *ppl* degrades phage and host DNA upon infection, but this process can be inhibited by gp130 (Fig. 4K).

Testing the phage triggers is challenging because reduced cell viability from phage gene toxicity and abortive infection, a common mechanism for anti-phage defense^71^, is phenotypically ambiguous. As a simple test to validate triggers, we expressed them on an arabinose-inducible plasmid and monitored cell growth in liquid culture after induction. Two of the six triggers identified by PhageMaP exhibited cell growth deficits when induced in the presence of a defense system but not in the control. Genes 40 and 123 had toxicity only in the presence of the *hhe* and Ec67 retron defense systems, respectively (Fig. 4L-M). In both cases, growth was impaired even in the repressed state, likely due to leaky expression (Fig. 4L-M). Overall, our data suggest that these genes are triggers for their respective defense systems. The absence of a cell growth defect in the remaining trigger-defense pairs can be explained by the following: (1) they may be false positives, (2) experimental conditions are not sufficient to induce abortive infection, or (3) they may not be abortive infection systems (Supplementary Fig. 7F).

In summary, our results provide proof of PhageMaP’s ability to identify key counters and triggers to anti-phage defense. We showed that the same phage genes can serve as counters for functionally distinct anti-phage systems, a trend also exemplified by the T7 Ocr protein with BREX and RM systems^51^. We demonstrated that by merging PhageMaP, structural information, and foundational knowledge from prior studies, mechanistic hypotheses can be generated. Because this panel is collectively represented in 32% of sequenced bacterial and archaeal genomes^33^, phage-antiviral defense interactions discovered may be generalizable to other phage-host pairs.

### PhageMaP identifies anti-phage defense counters and triggers in a model phage

We next sought to evaluate the fitness contributions of T7 genes during anti-phage defense. We obtained a total of >1900 T7 fitness measurements for a susceptible 23-member subset of the anti-phage defense panel. Due to the low correlation between replicates, BREX type I, Retron-TIR, and restriction-like systems were omitted from further analysis (Supplementary Fig. 6C). A total of 229 significant fitness scores (p < 0.05) were identified across 20 defense strains (Fig. 5A). Since the effect sizes in T7 (-1.9 to 1.8) were significantly lower than those in Bas63 (-5.0 to 2.5), we selected gene-defense pairs before multiple-testing correction to help capture subtle effects in our clonal validations. To better discriminate signal from noise, a minimum fitness score difference of 0.5 from the control was imposed, which resulted in a final set of 68 gene-defense pairs across 18 strains (Fig. 5A). Each defense system had distinct patterns of essentiality (Fig. 5A). Interestingly, seven genes (*g1.4*-*g2* and *g2.8*) found within the Class II T7 DNA metabolism cluster were identified to be potential triggers for *hhe* defense (Fig. 5A). Since it is unlikely that *hhe* is activated through direct recognition of each trigger, anti-phage activity is likely activated in response to the triggers’ functions rather than the triggers themselves (Fig. 5A).

**Figure 5.**
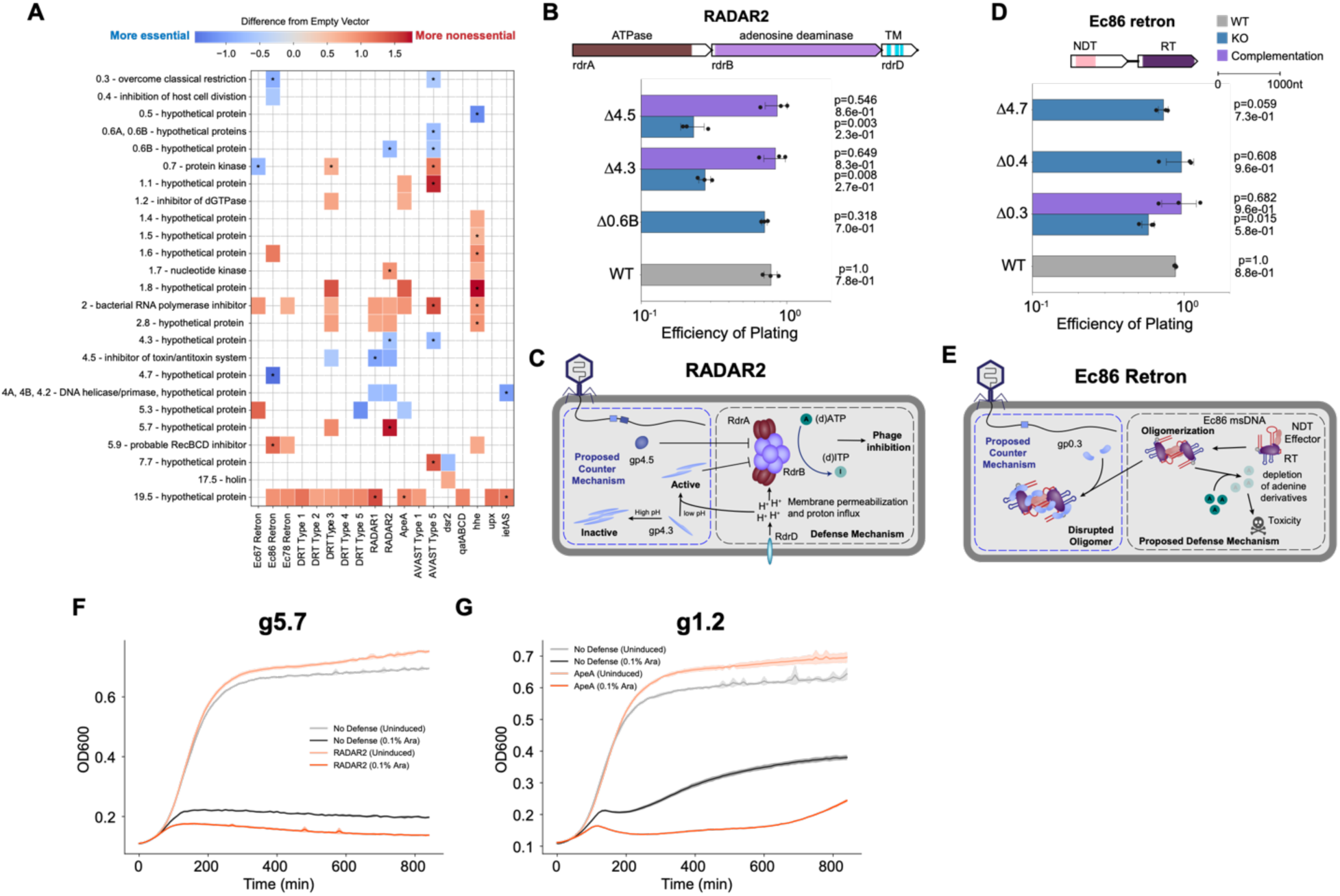
PhageMaP identifies anti-phage defense counters and triggers in T7 phage. **(A)** Plot showing all identified uncorrected significant hits (Welch’s t-test, p<0.05) and significant hits after multiple testing correction with the Benjamini-Hochberg method (q<0.1, denoted by an asterisk in each tile) with a score that is at least a difference of 0.5 from the control. The color of the tiles represents the difference in Z-normalized fitness scores between with defense and an Empty Vector control. **(B)** EOPs for T7 knockouts with the RADAR2 host. Average titers on the control strain were first determined (3 technical replicates). Each point on the plot represents a technical replicate for EOP using those calculated averages as the denominator for each titer measurement on the defense host. Bars represent the mean of three replicates, and error bars denote the standard deviation. Hypothesis testing was performed pairwise with each knockout (KO) and WT. P-values (Welch’s t-test) and means are annotated above the bars. Complementation host contained the defense system and a plasmid expressing deleted gene. **(C)** Proposed model of defense and counter-defense for RADAR2. **(D)** EOPs for T7 knockouts with the Ec86 retron host. Plot was generated the same as B. **(E)** Proposed model of defense and counter-defense for Ec86 retron. **(F)** Growth curves for expression of g5.7 in cells containing no exogenous defense or the RADAR2 system. 1 x 10^7^ cells were either repressed with 0.2% glucose (uninduced) or induced with 0.1% L-arabinose and grown for ∼14hrs. Central line represents the average of 3 replicates and the shaded flanking regions represent the standard deviation. **(G)** Growth curves for expression of g1.2 in cells containing no exogenous defense or the ApeA system. Experiment was performed and plot made the same way as F.

We tested 17 pairs with plating assays to validate counters, and 46 pairs were tested using cell growth assays to validate triggers (Fig. 5A). For the 17 counters, we compared the plating efficiency of T7 gene knockouts to wildtype phage. We expect that the removal of these potential inhibitors of anti-phage defense would reduce phage fitness. The average plating efficiency compared to wildtype only reached statistical significance in 3 out of the 17 gene-defense pairs tested (Fig. 5B and D, Supplementary Fig. 8A). The reduction in efficiencies of plating was never greater than 10-fold, indicating a correlation between plating efficiency and PhageMaP scores. The weak effect suggests that these genes may serve minor or supporting roles in the evasion of anti-phage activity.

For the *Pluralibacter gergoviae* RADAR system (RADAR2), gp4.3 (hypothetical) and gp4.5 were predicted counters (Fig. 5A). Deletion of either gene reduces the plating efficiency by ∼70% which was restored upon complementation (Fig. 5B). RADAR2 mediates phage defense through ATP deamination and consists of three genes^33,90^: RdrA, which contains an ATPase domain that facilitates deamination; RdrB, the adenosine deaminase; and RdrD, an auxiliary protein with putative membrane pore-forming function^95^. Gp4.3 and gp4.5 share structural similarities with two inhibitors of F_1_-ATP synthase ATPase activity: IF1 and the ɛ subunit^96,97^ (Supplementary Fig. 8B). Since RADAR and ATP synthase have ATPase function, gp4.3 and gp4.5 may be general ATPase inhibitors. Gp4.5 is also a potential counter to RADAR1 from *Citrobacter rodentium*, but the effect was not significant in our plating experiments, possibly due to the weak RADAR1-gp4.5 interaction (Supplementary Fig. 8A). Since gp4.3 is more essential when rdrD is present (i.e. in RADAR2 but not RADAR1), gp4.3 essentiality may be driven by rdrD (Fig. 5A). IF1, a structural homolog to gp4.3, functions as an active homodimeric inhibitor for F1-ATPase activity at reduced pH^98^. Collectively, we propose a model for RADAR2 defense in which adenosine deamination is impaired by gp4.5 and active gp4.3. Gp4.3 can be activated after RdrD pore formation induces cytoplasmic acidification (Fig. 5C).

For Ec86 retron, *g0.3* was a predicted counter (Fig. 5A). Deletion of *g0.3* (*ocr*) significantly reduced the plaquing efficiency of T7 by ∼34%, which was rescued upon gene complementation (Fig. 5D). Genes 0.4 (inhibitor of host cell division) and 4.7 (hypothetical) which were also predicted counter did not reach significance in our clonal tests (Fig. 5D). Within Ec86 oligomers, the msDNA makes crucial contact with a positively charged patch in the enzymatic N-lobe of the effector^81^. We speculate that the negatively charged, DNA-mimicking ocr protein can weakly interact with the effector and interfere with effector-mediated cell death (Fig. 5E). Indeed, gp0.3 has binding activity with DNA-interacting enzymes such as the bacterial RNA polymerase to inhibit transcription^99^. These results highlight the versatility of DNA mimicry and gp0.3, which is a potent inhibitor of Type I RM systems and BREX^51^, in evasion of other antiviral defenses.

From the 46 potential triggers, there was reduced cell growth in only 2 defense strains (Fig. 5F and G). This failure rate can be attributed either to the toxic expression of most genes -- even in the absence of anti-phage defense -- or to the requirement that phage genes be expressed during infection to elicit an anti-phage response (Supplementary Fig. 9). Cell growth decreased relative to the control when gp5.7 is expressed alongside RADAR2 and when gp1.2 is expressed with ApeA (Fig. 5F and G), suggesting that these genes encode triggers to their respective anti-phage defense system. Gp5.7 is an inhibitor of σ^s^-dependent transcription in *E. coli*^100^. A prior study demonstrated that gp5.7 inhibition of host transcription triggered nucleotide depletion through dCTP deamination and dGTP hydrolysis^101^. We show that RADAR2-mediated adenosine deamination is also activated by gp5.7. As an inhibitor of the dGTPase, gp1.2 serves a role in maximizing reserves of dGTP for phage DNA replication^102^. ApeA encodes a single protein that contains an RNase domain, suggesting defense is executed on RNA substrates^103^. ApeA may be activated by sensing the nucleotide pool, but the mechanism by which conservation of cellular dGTP through gp1.2 activates the RNase remains unclear.

In summary, we showed that PhageMaP has the sensitivity to capture subtle effects. Because small effects are often overlooked, yet informative, examining these weaker phenotypes can help generate interesting hypotheses and mechanistic insights. We ascribed new functions to genes with previously unknown (g4.3) and known (g0.3 and g4.5) roles.

### Transfer of a newly identified counters enhances infectivity of a recipient phage

To test if counters identified from one phage can be used to boost the infectivity of the other ineffective phage, we first examined if there is a defense strain with an identified counter and is fully susceptible to only either Bas63 or T7. Gp130 provides Bas63 with *ppl* immunity, a strain which has resistance to T7 (Fig. 4J). We find that *ppl* impaired plaque formation for T7 compared to the No-defense control (Fig. 6A-C). We tested if this phenotype is reversed by expression of gp130 from a plasmid or directly from the phage genome by replacing the nonessential *g4.*3 with gp130. Expression of gp130 but not a GFP control from a plasmid restored the T7 plaquing phenotype to that of the No-defense control (Fig. 6A). Interestingly, expression of gp130 from a plasmid improved plaquing efficiencies past the efficiencies seen in the No-defense control, suggesting gp130 may have a general role in robust phage growth (Fig. 6B). Expression of gp130 directly from the T7 genome improved plaquing efficiency ∼3-fold on the *ppl* strain compared to wildtype (Fig. 6A-B). Wildtype T7 forms small, turbid plaques on the *ppl* strain (Fig. 6C). Expression of gp130 either from a plasmid or genome resulted in large, clear plaques like the No-defense control (Fig. 6C). These data indicate that counters are modular and transferable and demonstrate how PhageMaP datasets involving bacterial immune systems can be leveraged to engineer more efficacious phages.

**Figure 6.**
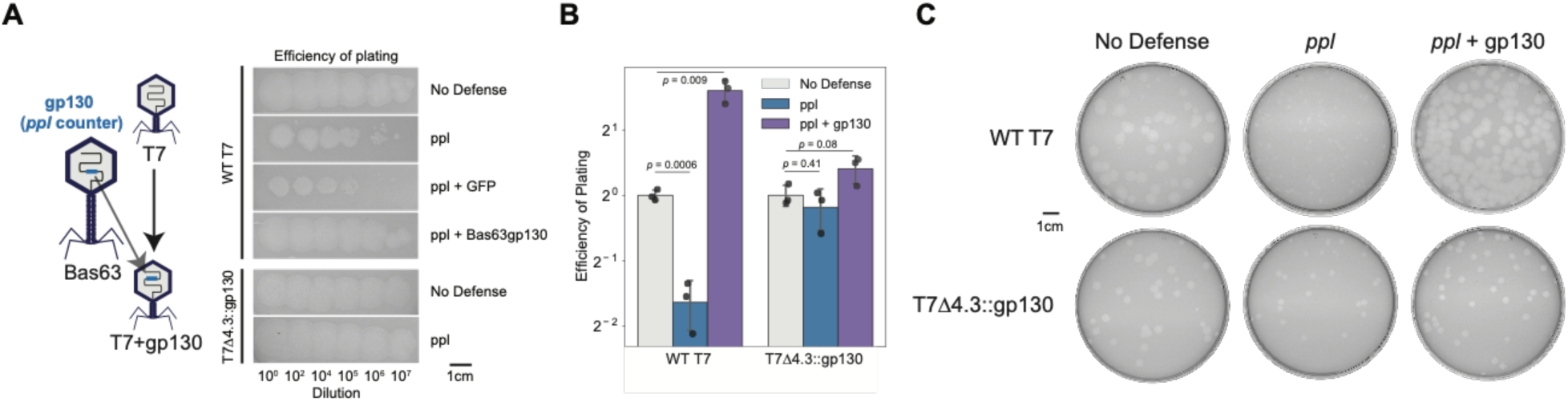
Transfer of gp130 from Bas63 to T7 improves infectivity on *ppl*. **(A)** Plaquing phenotypes of T7 phage with No-defense (empty vector), *ppl*, *ppl* + GFP, and *ppl* + Bas63gp130 hosts and of T7Δ4.3::gp130 on No-defense and ppl hosts. **(B)** Efficiency of plating (EOP) of T7 phage and T7Δ4.3::gp130 on No-defense, *ppl*, and *ppl* + Bas63gp130 hosts. For each phage, the No-defense is used as the reference for EOP calculations. Error bar represents standard deviation of three replicates. P-values are calculated using Welch’s t-test. **(C)** Plaque sizes of T7 phage and T7Δ4.3::gp130 on No-defense, *ppl*, and *ppl* + Bas63gp130 hosts. Images are representative of three replicates.

## Discussion

High-throughput genetic screens have revolutionized the way we study phage-host interactions and discover gene function^19,29,30,104–109^. PhageMaP leverages the pooled barcoding of genome-wide loss-of-function phage mutants to systematically and efficiently interrogate conditional essentiality in high-throughput. Using PhageMaP, we presented the first comprehensive conditional gene essentiality maps for model phage T7 and non-model phage Bas63 in the context of laboratory and anti-phage defense strains. From these datasets, we found large portions of the phage genomes are nonessential (>50% of genes) in the tested conditions and identified novel counters and triggers with varying sensitivities for several anti-phage defense systems. We demonstrated that phage genes, such as *g79* and *g80* in Bas63 and *g0.3* and *g4.5* in T7, can counter multiple distinct defense systems, underscoring the versatility of the toolbox found in phage genomes. PhageMaP also captured co-essential genes that function in the same pathway (e.g. *g79* and *g80* in Sir2-HerA defense and *g84* and *g85* in *qatABCD*). We also showed a counter (gp130) from one phage (Bas63) can be used to increase infectivity of another (T7) in the presence of an anti-phage defense (*hhe*). By combining PhageMaP results, sequence and structural alignment tools, and prior literature, we proposed mechanistic models for previously unknown phage-host interactions to encourage future studies. PhageMaP can be coupled with bioinformatic approaches to characterize analogous genes across diverse phages, thereby accelerating functional annotation of unknown viral sequences.

PhageMaP currently has some limitations. First, since single knockouts are generally made, genes that are functionally redundant within a phage genome may be classified as nonessential despite collectively playing roles in essential processes. Second, we have only demonstrated generalizability to dsDNA coliphages. While application to non-*E. coli* systems is highly feasible, the recombineering host must be able to both house plasmid libraries and express Cas proteins and recombination machinery. Because plasmid vectors for non-*E.* coli strains are available, and CRISPR-Cas systems and homologous recombination are ubiquitously utilized in microbes^34^, PhageMaP can be ported over to non-*E. coli* systems. Nonetheless, our application using coliphages still reveals general strategies for anti-phage defense and counter-defense as seen with RADAR2, a non-*E. coli* defense system. Third, phages resistant to genetic manipulation through DNA modification or protection may require prior knockout of the resistance factor or re-engineering of the Cas proteins or recombination machinery^37,110^.

Phages must carry essential core genes and auxiliary genes, such as those for evading host defense. Although it might seem that phages could continuously acquire adaptations to enhance their fitness in any environment, several trade-offs limit this. Larger genomes impose greater energetic demands and can reduce virion stability^3^. Furthermore, an increased genome size after acquisition of one antiviral inhibitor may sensitize the phage to another anti-phage defense system (e.g. restriction-modification or CRISPR-Cas) triggered by the newly obtained elements^111^. How phages evolved to balance adaptability and genome size is an open question. The versatility of counters to different anti-phage systems demonstrated in our study suggests that phages evolved multi-purpose proteins as a solution to these trade-offs. Such proteins provide additional functionality without compromising genomic constraints. This strategy is also exemplified in the straight tail fiber protein of coliphage T5 which dually functions as a tape measure protein and as an enzyme that degrades the cell wall and fuses the host membranes^112^. With further PhageMaP screening, we anticipate uncovering more hidden functions in already annotated proteins.

PhageMaP opens the door to new research directions. In this study, we applied PhageMaP to a small representative subset of possible anti-phage defense systems found in nature, but new systems are constantly being discovered which provides a continuous stream of opportunities to investigate phage-defense interactions with PhageMaP^33,71^. PhageMaP can also be extended to study how genetic factors affect phage infectivity in different environmental, growth, or media conditions^113–117^. Because PhageMaP is host-agnostic, it can be applied to complex microbial communities to uncover differential gene essentiality dependent on microbiota composition^118^. The PhageMaP datasets have exciting applications in rational phage engineering. The identification of broadly nonessential genes facilitates the construction of reduced phage genomes which can overcome packaging limits^18^ and serve as scaffolds to introduce non-native genes that boost phage efficacy^22,119^. These beneficial non-native genes can be discovered through PhageMaP screening. For example, anti-phage counters from one phage can be added, and triggers can be removed from the phage genome to improve infectivity on a target host carrying certain defense systems. We anticipate that with the PhageMaP testing of additional phages, especially jumbo phages^120^, many more interesting facets of phage biology can be uncovered. PhageMaP is positioned as a discovery tool for identification of novel phage-host interactions and functional characterization of genes.

## Methods

### Strains and culture conditions

Bacteriophage T7 was obtained from ATCC (ATCC BAA-1025-B2). Bacteriophage Bas63 (JohannRWettstein) was obtained from Alexander Harms (Biozentrum, University of BaselBiozentrum) and derived from the BASEL collection^32^. *E. coli* DH10B (C3020) and *E. coli* BL21(DE3) (C2527) were purchased from NEB (C3020). *E. coli* BL21 (REL606) was obtained from Robert Landick (University of Wisconsin-Madison). *E. coli* S17-1 was obtained from Andrew Hryckowian (University of Wisconsin-Madison). *E. coli* BW25113 was obtained from Douglas Weibel (University of Wisconsin-Madison) and derived from the Keio collection^13^. *E. coli* LE392 (bor::kanR) was obtained from Lanying Zeng (Texas A&M University). *E. coli* UTI33 and UTI46 were obtained from Rod Welch (University of Wisconsin-Madison) and derived from the UTI collection^42^. The *E. coli* ECOR13 and ECOR4 strains were obtained from the Michigan State University STEC Center and derived from the ECOR collection^43^. *E. coli* MG1655 is a laboratory strain. The phage defense strains in *E. coli* DH5a were a gift from Feng Zhang (Addgene plasmids #157880-157912)^33^ and the defense plasmids were subsequently transformed into DH10B for T7 selection. Bas63 selection used the strain background (NEB Stable) provided by Addgene. For long term storage, all bacterial strains were stored at -80°C in 25% glycerol and 75% LB.

All bacterial strains were grown in LB media (1% tryptone, 0.5% yeast extract, 1% NaCl, and 1.5% agar for plates or 0.7% agar for top agar). If applicable, antibiotics kanamycin (Kan 50µg/mL final concentration), chloramphenicol (Cam 25µg/mL), and spectinomycin (Spec 100µg/mL final concentration) were added for plasmid maintenance. Inducers anhydrotetracycline (1µM final concentration) and L-arabinose (0.1%) were added if necessary for induction of pBAD promoter. D-glucose (0.2% or 0.5% final concentration) was added if necessary for repression of pBAD promoter. All incubations were performed at 37°C and shaken at 200-250rpm, if growing liquid cultures, unless specified otherwise.

The initial laboratory stock of bacteriophage T7 was propagated using *E. coli* BL21 after receipt from ATCC. New stocks were made by propagating the initial laboratory stock with *E. coli* DH10B. Bas63 was also propagated on DH10B. Propagation of both phages was performed using LB and culture condition described above. Phages were purified with 0.22µm filters and stored at 4°C in LB.

*E. coli* competent cells were prepared by inoculating 5mL of overnight culture into 400mL of SOB media (2% tryptone, 0.5% yeast extract, 10mM NaCl, 2.5mM KCl, 10mM MgCl_2_, 10mM MgSO_4_). Cells were grown at 37°C shaking at 200rpm until OD_600_ ∼0.6 as determined by an Agilent Cary 60 UV-Vis Spectrometer and Ultrospec 10 Cell Density Meter (Amersham Biosciences). Cells were spun down at 5.5k x g for 10 minutes at 4°C.Two washes with cold 10% glycerol were performed. Cells were spun down at 5.5k x g for 10 minutes at 4°C after each wash, and the supernatant was removed. The pellet was then resuspended with 1:100 starting volumes of cold 10% glycerol. Cells were aliquoted and stored at -80°C. Alternatively, chemically competent cells were made using the Mix & Go! *E. coli* Transformation Kit and Buffer Set (Zymo Research #T3002) according to manufacturer instructions.

An estimate of 8 x 10^8^ CFU/mL at OD_600_ 1 was used for quantifying cell numbers in a volume of culture in all experiments. Cell numbers were used to calculate amount of phage to add for a target multiplicity of infection (MOI).

### Double agar overlay plaque assays

Plaque assays were routinely performed to isolate phages and quantify titer of phage preparations. In general, 200-300µL of stationary phase target host and a dilution of phage were added to 4mL of 0.5-0.7% top agar (1% tryptone, 0.5% yeast extract, 1% NaCl, and 0.5-0.7% agar) with additional antibiotics, inducer, or glucose if applicable. The mix was then vortexed and plated on LB plates. The top agar was allowed to solidify (∼10 minutes) before incubation at 37°C overnight. Plates containing 10 to 1000 plaques were typically counted. Titers were calculated by averaging at least 2 replicates.

### General cloning procedures and base plasmid construction

All PCR reactions were performed using KAPA HIFI (Roche #KK2101) according to manufacturer instructions and with the following cycle settings unless specified otherwise: 95°C 3 minutes, (98°C 20 seconds→65°C 15 seconds→72°C X seconds) x 25 cycles, 72°C X seconds, and 4°C hold, where X varies depending on amplicon length. An initial denaturation step of 5 min at 95°C is used if the template is phage. A 2X KAPA HIFI master mix was made by adding dNTPs, HF Buffer, and KAPA HIFI DNA Polymerase and stored at -20°C. All PCR reactions use this master mix. DpnI (NEB #R0176L) digestion was performed if the template is derived from plasmid DNA by adding 1µL DpnI and 2.3uL 10X Cutsmart Buffer directly into the PCR sample, incubating at 37°C for 1 hour, and heat inactivating for 20 minutes at 80°C. DNA purification after PCR was performed either with the E.Z.N.A Cycle Pure Kit (Omega Bio-tek #D6492-01) or with gel extraction from 1% agarose gels using the E.Z.N.A Gel Extraction Kit (Omega Bio-tek D2500-01), both with a centrifugation protocol according to manufacturer instructions.

Golden Gate Assemblies were performed using the New England Biosciences (NEB) Golden Gate Assembly BsaI-HFv2 (NEB #E1601L) or BsmBIv2 (NEB #E1601L) Kits. Reactions were prepared according to manufacturer instructions, but cycling was performed for 60 total cycles with 5-minute steps. Gibson Assemblies were performed using either in-house Gibson Master Mix (final concentration 100mM Tris-HCl pH 7.5, 20mM MgCl_2_, 0.2mM dNTP, 10mM DTT, 5% PEG-8000, 1mM NAD+, 4U/mL T5 exonuclease, 4U/uL Taq DNA ligase, 25U/mL Phusion polymerase) or NEB Gibson Assembly Master Mix (NEB #E2611), following the Gibson Assembly Protocol (NEB #E5510).

Following assembly of constructs, reactions were diluted 5-fold with dH_2_O if performing plasmid transformation or dialyzed using MF-Millipore 0.025µm MCE membranes (Millipore Sigma # VSWP02500) in dH2O for 1 hour if performing donor library transformation or rebooting phage. For single plasmid transformations, 2µL of the dilution was transformed into 25µL DH10B electrocompetent cells. For library transformations, 5µL of dialyzed reaction was transformed into 25µL DH10B electrocompetent cells. For rebooting phage, half of each dialyzed reaction (∼10µL) is transformed into 50µL DH10B electrocompetent cells. Transformations were performed using a Bio-rad MicroPulser (#165-2100). For single plasmid and library transformations, the Ec1 setting (1-mm cuvette, 1.8kV) was used. For rebooting phage, the Ec3 setting (2-mm cuvette, 3kV) was used. Recovery after electroporation was performed in a shaker at 200-250rpm in a total volume of 1mL SOC (2% tryptone, 0.5% yeast extract, 10mM NaCl, 2.5mM KCl, 10mM MgCl_2_, 10mM MgSO_4_, 20mM glucose) for 30-60min. For phage reboots, this step was omitted and 300µL of transformants was added to 0.5% top agar and plated on LB plates. For library and single plasmid transformations, a diluted recovery was plated on LB plate with relevant antibiotics. Single colonies were picked and validated with PCR and/or sequencing prior to further outgrowth. Plasmid extraction was performed using the ZR Plasmid Miniprep – Classic Kit (Zymo Research #D4016) according to manufacturer instructions.

DNA quantification was performed using NanoDrop 2000 (Thermo Scientific) with 1.5 µl of DNA, except for Next Generation Sequencing DNA, which was quantified using a Qubit 4 fluorometer (Thermo Scientific) with the Qubit 1X dsDNA HS Assay Kit (Thermo Scientific, #Q33231). Following the manufacturer’s documentation.

Following construction of non-library plasmids, plasmids were verified with either Sanger Sequencing (Functional Bioscience) or whole plasmid sequencing (Plasmidsaurus).

#### Construction of pUC19-Donor-BsmBI

To generate pUC19-Donor-BsmBI, the landing vector for oligos encoding donor and sgRNA information, two two-part Gibson Assembly reactions were performed. In the first reaction, one fragment contains the pUC19 vector with the lacZa reporter gene removed, and the second fragment contains donor DNA and the sgRNA for one locus in T7. For the second Gibson reaction, one fragment is an amplicon that contains the sgRNA scaffold with the pUC19 vector generated from the first reaction. The second fragment contains an sfGFP gene flanked by new BsmBI cutsites and sequences homologous to the first two fragments. This version of the landing vector contains an ampicillin resistance cassette. This cassette was then swapped with a kanamycin cassette from a previously synthesized laboratory plasmid. Site-directed mutagenesis through Gibson Assembly was done using this plasmid to remove the native BsaI recognition sites and generate the final pUC19-Donor-BsmBI plasmid. A variant of this plasmid (*pUC19-Donor-BsaI)* containing BsaI sites was also constructed using site-directed mutagenesis of pUC19-Donor-BsmBI and a 2-part Gibson Assembly.

#### Construction of pCas9-RecA/sfGFP

A three-part Gibson Assembly was used to generate tetracycline-inducible pCas9-RecA and pCas9-sfGFP. First the plasmid vector containing an SC101 origin, kanamycin resistance cassette, and the tet expression system was amplified with PCR from a previously generated laboratory plasmid. To obtain the Cas9 insert, a laboratory plasmid SC101-Cas9 was used as template for PCR. The recA insert was extracted from the pMP11 plasmid obtained from Brian Pfleger (University of Wisconsin-Madison). The sfGFP insert was extracted from the pUC19- Donor-Template plasmid.

#### pSM103 expression constructs

pSM103, which contains an arabinose-inducible pBAD promoter driving sfGFP, a kanamycin resistance cassette, and a pET31b origin of replication, was previously generated in the lab and used as the vector for insertion of phage genes for complementation and trigger studies. The vector was amplified with PCR. Phage genes were PCR amplified from 1uL of phage stocks (∼10^10^ PFU/mL) and contain homology to the termini in the amplified pSM103 vector. The resulting fragments were assembled with Gibson Assembly.

### Donor plasmid library design and construction

#### Donor sequence design

The donor plasmid inserts can be found in Table S1. T7 (V01146) and Bas63 (MZ501086) genome files were obtained from GenBank. To design sgRNAs for T7, DeepSpCas9 and CRISPRon gRNA efficiency prediction tools were used^122,123^. We find that the predicted efficiencies were concordant and opted to only use CRISPRon to design the Bas63 sgRNAs. For each sgRNA, the theoretical cut site (3nt upstream of the PAM) was used as the starting position for the design of the 95nt homology arms. The homology arms are designed such that the barcode will be inserted at the cut site position. The sgRNA, 5’ homology arm, 3’ homology arm, J23119 promoter, primer binding, and barcode landing sequences were combined with a custom Python script to create the donor plasmid insert sequences. These sequences were ordered as oligo pools (Twist Bioscience).

#### Construction of the barcode insert

The barcode insert contains a 16N sequence flanked by two constant sites containing stop codons in every reading frame and is used as primer binding sites to extract barcodes for sequencing. At the termini of the insert are BsmBI or BsaI cutsites for assembly into the donor plasmid vector. BC_oligo100, which contains the 16N barcodes, is first annealed and extended with either Twist_Bcoligo100_F (BsaI termini) or Twist_Bcoligo100_BsmBI_F (BsaI termini) by adding 1µL of each oligo (100µM) to 10µL 2X KAPA HIFI Master Mix and 8µL water. This mix was incubated at 95°C for 30 seconds, 63°C for 30 seconds, and then 72°C for 30 seconds. The annealed oligos were then diluted 10-fold with the addition of 180µL of water. 1µL of the dilution was used as template with primer pairs (10uM final concentration) Twist_Bcoligo100_F + Twist_Bcoligo100_R (BsaI termini) or Twist_Bcoligo100_BsmBI_F + Twist_Bcoligo100_BsmBI_R (BsmBI termini). PCR amplification was performed as follows: 95°C 3 minutes, (98°C 20 seconds→66°C 15 seconds→72°C 15 seconds) x 20 cycles, 72°C 15 seconds, and 4°C hold. To mitigate PCR bubble formation, one additional round of PCR was performed after the addition of 1uL of each primer. Each barcode insert was gel extracted and quantified as described above.

#### Insertion of the donor oligo sequences into pUC19 vector

To insert the T7 donor oligo sequences into the pUC19-Donor-BsmBI vector, a Golden Gate Assembly (BsmBI) was performed using 75ng of pUC19-Donor-BsmBI and a 2-fold molar ratio of oligo insert. For Bas63 donor oligo sequences, pUC19-Donor-BsaI vector was used with Golden Gate Assembly (BsaI) Mix. The reactions were performed as described above. The reactions were dialyzed and transformed as describe above. Recovery was performed in SOC for 30 minutes. 100µL of the recovered transformants was plated on LB-Kan plates and the rest was used to inoculate 5mL of LB-Kan media for overnight the growth. Plasmid DNA from the resulting outgrowth was extracted as described above. The resulting purified plasmids pUC19- T7Library and pUC19-Bas63Library are the landing vectors for the barcode insert.

#### Insertion of the barcode into donor vector

To insert the barcode insert (BsaI termini) into pUC19-T7Library, a Golden Gate Assembly (BsaI) was performed using 75ng of pUC19-T7Library and a 2-fold molar ratio of barcode insert.

For barcode insert (BsmBI termini), pUC19-Bas63Library vector was used with Golden Gate Assembly (BsmBI) Mix. The dialysis, transformation, and recovery outgrowth were handled the same as during the donor oligo sequence insertion. Serial dilutions of the recovered transformants were plated on LB-Kan plates and used to inoculate 5mL of LB-Kan media for overnight the growth. We proceeded with the dilution that gave approximately 10-fold coverage of the theoretical library size. Plasmid DNA from the resulting outgrowth was extracted as described above. The resulting plasmid libraries are pUC19-T7LibraryBC and pUC19- Bas63LibraryBC.

#### Transformation of pUC19-T7LibraryBC and pUC19-Bas63LibraryBC into Cas9-RecA cells

*E. coli* DH10B Cas9-RecA competent cells were made the same way as described with *E. coli* DH10B with three differences. Cultures were scaled down to 10mL, all growth steps occurred at 30°C, and the LB media used was supplemented with spectinomycin. 1ng of pUC19- T7LibraryBC and pUC19-Bas63LibraryBC was transformed into 25µL of Cas9-RecA competent cells. Transformants were recovered at 30°C for 30 minutes. Serial dilutions of the recovered transformants were plated on LB-Kan-Spec plates and used to inoculate 5mL of LB-Kan-Spec media for overnight the growth. We proceeded with the dilution that gave approximately 10-fold coverage of the theoretical library size. Glycerol stocks for both libraries were made and stored at 80°C. Plasmid DNA from the resulting outgrowth was extracted as described above.

#### Sequencing pUC19-T7LibraryBC and pUC19-Bas63LibraryBC

To generate sequencing amplicons for both libraries, 1ng of plasmid DNA was used as template for a 10µL PCR reaction with primer pairs DelLib{2,3,4}N_NGS_F + DelLib{2,3,4}N_NGS_R containing 2, 3, and 4 N offsets and the following PCR cycle settings: 95°C 5 minutes, (98°C 20 seconds→65°C 15 seconds→72°C 15 seconds) x 15 cycles, 72°C 1 minute, and 4°C hold.

Illumina i7 and i5 adapter sequences containing unique indices and were added after performing a second 25µL PCR reaction with the following cycle settings: 95°C 3 minutes, (98°C 20 seconds→65°C 15 seconds→72°C 15 seconds) x 10 cycles, 72°C 1 minute, and 4°C hold.

PCR products were purified with the the E.Z.N.A Cycle Pure Kit and quantified using the Qubit 4 as described above. The purified products were pooled and sequenced on an Illumina MiSeq with a 2 x 250 Miseq Reagent v2 kit.

#### Sequencing data processing and analysis

Read pairs were merged and filtered with Fastp v0.23.4^124^, requiring a minimum of 90% of the read having scores above Q25. The adapter sequences were removed, and relevant donor sequences were extracted using Cutadapt v4.1^125^. The resulting sequences were further analyzed using custom Python scripts. The result is the mapping of barcodes to their theoretical insertion site (genic or intergenic regions).

### Generating PhageMaP phage libraries

#### Homologous recombination with donor libraries

For both T7 and Bas63 PhageMaP libraries, 5mL of LB-Kan-Spec was inoculated with 100µL of pUC19-Bas63Library + pCas9-RecA or pUC19-T7Library + pCas9-RecA thawed glycerol stock. Both cultures were grown overnight at 30°C. The T7 donor library was added to 800mL of LB-Kan-Spec, and the Bas63 donor library was added to 500mL of LB-Kan-Spec. Both were grown at 30°C until the OD_600_ reached 0.3-0.5. Anhydrotetracycline was added to each culture to a final concentration of 1µM to induce Cas9 and RecA. Cultures were grown at 30°C until the OD_600_ reached ∼0.6. WT phage was added to their respective libraries at an MOI of 5. We reasoned that infection of the recombination host at a high multiplicity of infection (MOI 5) would enable phage variants with knockout of essential genes to persist in the population due to complementation from secondary co-infections. After infection of the donor host with wildtype phage, the genomes are inactivated by the encoded Cas9-sgRNA pair to create double-stranded breaks used for HR and reduction of wildtype background. Inactivated phage genomes undergo RecA-mediated HR with the intracellular donor plasmids to restore genome integrity and to insert barcodes into target locus. After a 2-hour incubation at 37°C, lysates were transferred to centrifuge bottles and centrifuged at 16,000 x g for 15 minutes at 4°C to remove cell debris. The supernatant was then filtered using 0.45µm PES membrane filters.

#### Cesium Chloride (CsCl) density gradient

CsCl density gradients were used to concentrate and purify phage libraries. To remove free nucleic acids, lysates were first treated with DNase I (Roche #10104159001, 1µg/mL final concentration) and RNase A (Fisher Scientific # NC972993, 10µg/mL) after the addition of CaCl_2_ (500µM) and MgCl_2_ (2.5mM). Treatment was performed for 1.5 hour at room temperature with gentle stirring. NaCl was then added to a final concentration of 1M and allowed to dissolve.

Next, 80g of PEG-8000 was added to the lysate and stirred at room temperature until dissolved. The lysates were then stirred at 4°C for 45 minutes and subsequently incubated overnight at 4°C without stirring for precipitation. The precipitated lysates were centrifuged for 10,000 x g for 15 minutes at 4°C. The supernatant was removed, and the pellets were allowed to dry at room temperature. Pellets were resuspended in 7mL of SM Buffer (100 mM NaCl, 25 mM Tris-HCl pH 7.5, 8 mM MgSO_4_). To remove the insoluble material, the resuspended pellets were centrifuged at 3000 x g for 10 minutes twice, moving the supernatant to a new tube after each spin. CsCl was added to the supernatant at a rate of 0.75g/mL. The phage library solutions were added to a step gradient of 1.4, 1.5, and 1.6g/mL CsCl in 14mL Ultra-Clear centrifuge tubes (Beckman Coulter #344060) and centrifuged in a SW40 Ti rotor (Beckman Coulter) at 24,000 x rpm for 24 hours at 5°C. For each tube, a faint light blue band was observed between the 1.4 and 1.5 layers and extracted using a syringe and a 26-gauge needle. The purified phage libraries were then dialyzed overnight at 4°C using Slide-A-Lyzer cassettes with a 10k MWCO (Thermo Scientific #66380) in SM Buffer supplemented with 1M NaCl. Two additional rounds of dialysis were performed in normal SM Buffer for 2 hours at room temperature. After dialysis, the phages were extracted from the cassettes and filtered with a 0.22µm PES membrane filter. Plaque assays (described above) using DH10B as a host were performed to titer the libraries.

#### Sequencing PhageMaP libraries

Phage genomic DNA from libraries were extracted using the Norgen Phage DNA Isolation Kit (Norgen #46800) with DNase and proteinase K treatment, following manufacturer instructions. Primers BC_2N_Twist_NGS_F and BC_4N_Twist_NGS_R were used to amplify barcodes for sequencing using the following PCR cycle settings in 20µL reactions: 95°C 3 minutes, (98°C 20 seconds→65°C 15 seconds→72°C 15 seconds) x 20 cycles, 72°C 1 minute, and 4°C hold.

Illumina i7 and i5 adapter sequences containing unique indices and were added after performing a second 25µL PCR reaction with the following cycle settings: 95°C 3 minutes, (98°C 20 seconds→65°C 15 seconds→72°C 15 seconds) x 15 cycles, 72°C 1 minute, and 4°C hold.

Libraries were quantified using a Qubit 4 as described above. The T7 library was sequenced using a 2x150 Miseq Nano Kit (Illumina). The Bas63 library was sequenced using a 2x250 Miseq Nano Kit (Illumina).

#### Sequencing data processing and analysis

Read pairs were merged and filtered with Fastp v0.23.4^124^, requiring a minimum of 90% of the read having scores above Q25. The adapter sequences were removed, and barcodes sequences were extracted using Cutadapt v4.1^125^. The resulting sequences were further analyzed using custom Python scripts.

#### ddPCR for recombination efficiency estimate

Digital droplet PCR was performed to estimate abundance of recombinant population in the phage library as described in Supplementary Figure 2D. The estimation was calculated by first finding the percentage of phages that contain a barcode and then finding the percentage of barcodes that belong to the residual donor plasmid after phage library purification. The percentage of phages that contain a barcode was determined with a primer-probe pair where one primer-probe produces signal for the barcode insert (present in donor plasmids and recombined phage) and another primer-probe produces signal for a constant region in the phage (present in both recombined and unrecombined phage). The ratio of the former to the latter is the amount of barcode containing DNA detected relative to phage DNA. To account for the donor plasmid population, a second primer-probe pair is used where one primer-probe produces a signal for the kanamycin cassette found in the donor plasmid and another primer-probe produces a signal for the barcode insert (present in donor plasmids and recombined phage). The ratio of the former to the latter gives an estimate of the percentage of barcodes that originate from plasmid DNA. Multiplying the first ratio with the difference between 1 and the second ratio gives us an estimate of the percentage of phages with barcodes (recombinant percentage). For T7 libraries, 3pg of DNA was used for each reaction. For Bas63, 6pg of DNA was used. All reactions were performed in 20µL, using ddPCR supermix for probes (no dUTP) (Biorad #1863024), 0.9µM of each primer, and 0.25µM of each 5’ 6-FAM/ZEN/3’ IBFQ or 5’ HEX/ZEN/3’ IBFQ probe with the QX200 Droplet Digital PCR System (Biorad). Reactions were performed using the following PCR cycle settings: 95°C 10 minutes, (94°C 30 seconds→60°C 1 minute) x 40 cycles, 98°C 10 minute, and 12°C hold. Data were collected using the QX Software (Biorad) and further processed with custom Python scripts.

### PhageMaP selection experiments and analysis

#### PhageMaP selection

Overnight cultures of the target hosts were back diluted 1:10 in LB with additional supplements if necessary. Cultures were grown until mid-late log phase (OD_600_ ∼0.7-1) and aliquoted into three separate tubes to serve as replicates. The phage library was added at an MOI 0.1 to ensure comprehensive assaying of the library and single infections. All cultures were allowed to lyse at 37°C. For the laboratory *E. coli* strains infected by T7, cultures were collected after 1.5 hour. For the phage defense strains infected by T7, cultures which had visual lysis were collected after 3 hours. For the laboratory *E. coli* strains infected by Bas63, cultures were allowed to lyse overnight. For the 9 phage defense strains, including empty vector, infected by Bas63, cultures were allowed to lyse for 7 hours. All lysates were purified using 0.22µm PES membrane filters.

#### Sequencing phages after selection

Three pairs of primers containing 1, 2, or 3 N offsets and a unique 3 nucleotide barcode (for demultiplexing of replicate numbers) were used to amplify barcodes from 10µL of phages after selection with the following PCR cycle settings in 25µL reactions: 95°C 5 minutes, (98°C 20 seconds→65°C 15 seconds→72°C 15 seconds) x 20 cycles, 72°C 1 minute, and 4°C hold.

Illumina i7 and i5 adapter sequences containing unique indices and were added after performing a second 25µL PCR reaction with the following cycle settings: 95°C 3 minutes, (98°C 20 seconds→65°C 15 seconds→72°C 15 seconds) x 15 cycles, 72°C 1 minute, and 4°C hold.

Libraries were quantified using a Qubit 4 as described above. The T7 selections were sequenced using 1x100 NextSeq P2 Kits (Illumina) on a NextSeq 1000. The Bas63 selections were sequenced using a 2x150 NovaSeq Kit (Illumina) through the University of Wisconsin Biotechnology Center Sequencing Core.

#### Analysis of selection data

Read1 in each sequencing run was filtered with Fastp v0.23.4^124^, requiring a minimum of 90% of the read having scores above Q25. If this criterion was too stringent and most reads were discarded, a more lenient 80% of the read having scores above Q20 was imposed. The adapter sequences were removed, and barcode sequences were extracted using Cutadapt v4.1^125^. The resulting sequences were further analyzed using custom Python scripts. Barcodes mapping to the same sgRNA were summed. sgRNAs which do not have at least 2 reads post-selection and 15 reads pre-selection were excluded from analysis. Calculation of fitness scores were adapted from Enrich2^126^. Log ratios (*L)* for total count (c) of each sgRNA (*s)* in the initial (*i)* and selected *(sel)* were calculated using the following formula:

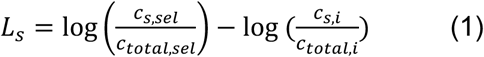

The standard error (SE_s_) for each log ratio was then calculated using the following formula:

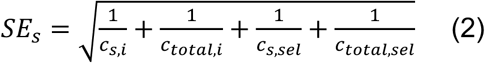

The log ratios and standard errors for all sgRNAs of the same gene were combined into a single gene score and standard error using the Enrich2 restricted maximum likelihood estimation with 50 Fisher scoring iterations. Overlapping regions of genes and intergenic regions are scored as individual genes. These genes scores from each of the replicates were once again combined into a single composite gene score using the same restricted maximum likelihood estimation. To help compare scores across conditions, each composite gene score was Z-normalized within each condition. To identify significant genes for the phage defense screens, these Z-normalized composite gene scores and their standard errors of phage defense conditions were compared to No-defense conditions using Welch’s two-sided t-test. A 0.5 minimum fitness score difference from control, pre-multiple testing correction significance threshold of 0.05, and post-correction threshold of 0.1 were set. For the anti-phage datasets, an essentiality threshold of 1.5 standard deviation (for T7) or 1 standard deviation (for Bas63) above the mean fitness score of known DNA replication and structural genes within each phage in the control/empty vector was set to enable binary labeling of each gene as essential or nonessential. Essential genes that became significantly more essential were omitted from further analysis as measurements were typically noisy due to the low initial counts and less likely to be biologically meaningful.

#### Additional analysis

PADLOC was used to identify the anti-phage defense systems found within a genome using available Genbank file inputs for each strain^121^. Structures of phage and defense proteins were predicted using the ColabFold (AlphaFold2) webserver with default settings^127,128^. Manual annotation of probable function in hypothetical genes was performed using BLASTp^129^. The most probable (E-value based), descriptive annotation at the time of search was used. DALI^130^ webserver was used to identify structurally similar proteins either within the full PDB database or AlphaFold Database v2 and to generate superimposed structures. PHASTEST was used to identify prophages within a genome using GenBank files^54^.

### Quantifying Cas9-RecA/sfGFP recombination efficiency

#### Construction of donor cells

Upon successful recombination, a 100bp deletion in T7g1.1 is replaced with 43bp containing a 11N barcode and barcode priming sites. A gene fragment containing 50nt homology arms, 11N barcode, J23119 promoter, sgRNA targeting T7g1.1, and BsmBI sites at the termini was PCR amplified using primers g576-Twist + 2969. Primers pUC19_BB_F + pUC19_BB_R were used to amplify the pUC19 vector with homologous ends to the gene fragment. These two fragments were assembled using Gibson Assembly and transformed using the transformation protocol outlined above. After plating, one colony was chosen after PCR and sequencing validation. This colony was used for miniprep. The resulting plasmid was transformed into *E. coli* DH10B Cas9- RecA or Cas9-sfGFP competent cells.

#### Testing recombination in Cas9-RecA and Cas9-sfGFP cells

Overnight cultures of Cas9-RecA and Cas9-sfGFP cells were back-diluted and grown to OD_600_ 0.3 at 30°C. To cells that require induction, anhydrotetracycline was added to a final concentration of 1µM. Uninduced and induced cells were grown for 1 hour at 30°C and subsequently diluted back down to OD_600_ 0.3. Wildtype T7 phage was added to 1mL of each culture in triplicate and allowed to infect for 1.5 hour. Lysates were spun down at 16,000 x g for 1 minute prior to filtering the supernatant using 0.22µm PES membranes.

#### Sequencing Cas9-RecA and Cas9-sfGFP lysates

To generate sequencing amplicons for both libraries, 1uL of phage lysate was used as template for a 25µL PCR reaction with primer pairs 576_NGS_4N_NGS_F + 576_NGS_4N_NGS_R containing 4N offsets and the following PCR cycle settings: 95°C 5 minutes, (98°C 20 seconds→60°C 15 seconds→72°C 15 seconds) x 12 cycles, 72°C 1 minute, and 4°C hold.

Illumina i7 and i5 adapter sequences containing unique indices and were added after performing a second 25µL PCR reaction with the following cycle settings: 95°C 3 minutes, (98°C 20 seconds→65°C 15 seconds→72°C 15 seconds) x 8 cycles, 72°C 1 minute, and 4°C hold.

PCR products were purified with the E.Z.N.A Cycle Pure Kit and quantified using the Qubit 4 as described above. The purified products were pooled and sequenced on an Illumina MiSeq with a 2 x 250 Miseq Reagent v2 kit.

#### Sequencing data processing and analysis

Read pairs were merged and filtered with Fastp v0.23.4^131^, requiring a minimum of 90% of the read having scores above Q25. The resulting sequences were further analyzed using custom Python scripts. Merged reads containing the known inserted barcode (’GTTTAGGCTGT’) and a consensus sequence (’GATTTAAATTAAAGAATTAC’) found in both recombined and unrecombined phages were quantified. Percent recombinant was calculated by finding the ratio of sequences containing the barcode and sequences containing the consensus sequence.

### Validation of individual hits from PhageMaP screening

#### Generating T7 knockouts

To determine if knockout of specific genes reduces the fitness of T7 relative to WT, efficiency of plating measurements were made between wildtype and knockout phages on relevant phage defense backgrounds. Amplicons were generated from the linear wildtype genome with homology at the termini to neighboring amplicons. Amplicons were assembled using a 5-part Gibson Assembly such that the central ∼80% of the target gene is deleted. 0.08 to 0.1pmol of each amplicon was used in each reaction. T7g0.3 knockout was previously constructed using a 6-part Gibson Assembly where a 173bp insert is inserted into position 212 of the gene.

Reactions were dialyzed and transformed into DH10B competent cells. After transformation, transformants were plated with 4mL top agar on LB plates and allowed to incubate at 37°C overnight. Single plaques were PCR verified before creating stocks using DH10B as the host.

#### Generating Bas63 knockouts

To determine if knockout of specific genes reduces the fitness of Bas63 relative to WT, efficiency of plating measurements were made between wildtype and knockout phages on relevant phage defense backgrounds. The knockouts were synthesized using Cas9-RecA-mediated homologous recombination and were designed such that the central ∼80% of the target gene is deleted. Gene fragments (Twist) containing 100nt homology arms, J23119 promoter, and sgRNA targeting the relevant position of the genome were inserted into pUC19-Donor-BsmBI with Golden Gate Assembly (BsmBI). Transformation and plating procedures were performed as described above. Single colonies were isolated, sequence verified, grown up for miniprep, and then transformed into DH10B Cas9-RecA competent cells. Colonies were grown overnight at 30°C. Overnight cultures of these strains containing donor plasmid and Cas9-RecA were back diluted 2-fold in LB-Kan-Spec and induced with anhydrotetracycline to a final concentration of 1µM for 30 minutes at 30°C. To perform recombination, 300µL of induced culture and 10^4^-10^5^ Bas63 phages were mixed into 0.7% top agar containing kanamycin, spectinomycin, and anhydrotetracycline. The top agar mix was poured onto LB plates. Plates were incubated at 37°C overnight. Successful plaques were verified with PCR prior to performing further experiments.

#### Efficiency of plating assays

To identify optimal dilutions to use for plaque assays, a spot plate assay was first performed using serial dilutions of the phages and spotting on a lawn of relevant hosts in 0.7% top agar.

Plaque assays were performed in triplicate by adding 300µL anti-phage defense host (or empty vector) and the optimal dilution of phage to 4mL of 0.7% top agar supplemented with chloramphenicol (25µg/mL) and pouring the mix into LB plates. For gene complementation hosts, kanamycin (50µg/mL) and 0.1% L-arabinose (w/v) was added to the top agar prior to pouring. Plates were counted after an overnight incubation at 37°C. Plates must have between 10-400 plaques to be included in further analysis. T7 genes 4.3 and 4.5 were too toxic when induced, thus complementation was performed from the leakiness of the uninduced cells.

#### Trigger assays

Candidate trigger genes were amplified from the wildtype genome via PCR. Amplicon ends contain homology to the vector backbone ends amplified from pSM103 with primers pSM103- BB-R + 1517. Constructs were assembled with Gibson Assembly as described above. The result is an arabinose-inducible expression plasmid for each candidate trigger gene. Plasmids were sequenced (Plasmidsaurus) prior to transformation into phage defense competent cells prepared the same way as described above. Transformants were recovered in SOC and plated on LB-Kan-Cam plates supplemented with 0.2% D-glucose (w/v) to repress expression of trigger genes. Individual clones were used to inoculate starter cultures supplemented with 0.2% D-glucose. 2 x 10^8^ cells from the overnight culture were spun down and resuspended in 100µL LB-Kan-Cam. 1 x 10^7^ cells (5µL) were added to 145µL of LB-Kan-Cam with either 0.1% L-arabinose (w/v) for induction or 0.2% glucose for repression on a 96-well plate in triplicate. OD_600_ was recorded every 10 minutes for 16 hours using a Synergy HTX Multi-Mode 96-well plate reader.

#### Competition Assay

Titers for barcoded WT Bas63 and knockouts were determined using plaque assays as described above. Overnight cultures of no defense, hhe, and hhe with gene complementation strains were back-diluted 1:10 and grown until OD 0.6-0.8. A 1:1 mix of the barcoded WT phage and knockout was added to each culture at an MOI of 0.01 in triplicate. The cultures were collected and filtered using a 0.22µm PES filter after ∼5hrs. 5uL of the purified lysate was added to fresh cultures for the second passage. The process was repeated for 3 passages. The initial mix, Passage 1, and Passage 3 lysates were used for sequencing on a 2x150bp NovaSeq run. Cutadapt v4.1 was used to extract barcodes and demultiplex replicates^125^. Futher analysis was done using custom Python scripts.

## Supporting information

phagemap_primers

suppdata1_T7_labecoli

suppdata2_Bas63_labecoli

suppdata3_Bas63_defense

suppdata4_T7_defense

## Acknowledgements

This work was supported by National Institute of Allergy and Infectious Diseases (NIAID) grant R21AI156785 (S.R.) and the NSF CAREER Award 2237251 (S.R.). J.C. was supported by the National Institute of General Medical Sciences of the National Institute of Health under Award Number T32GM135066 (Biotechnology Training Program). C.Y.M. and E.D.N. are supported by startup funds provided by the Department of Bacteriology at UW-Madison. We thank James Corban, Phil Huss, Nate Novy, Max Frenkel, Saylor Strugar, and Sarah Schmidt-Dannert for their feedback and helpful discussions in troubleshooting, data analysis, and phage biology. We thank Silas Miller for the plasmid vector used for some experiments and feedback on the manuscript. Some DNA amplicon sequencing was performed at the University of Wisconsin – Madison Biotechnology Center’s DNA Sequencing Facility (Research Resource Identifier – RRID:SCR_017759).

## Author contributions

Conceptualization, J.C and S.R.; Methodology, J.C.; Software, J.C.; Formal Analysis, J.C.; Investigation, J.C., E.D.N. C.C.; Resources, S.R. and C.Y.M.; Writing – Original Draft, J.C.; Writing – Review & Editing, J.C., E.D.N., C.Y.M., and S.R.; Supervision, S.R. and C.Y.M.; Funding Acquisition, S.R. and C.Y.M.

---

**Supplementary Figure 1.**
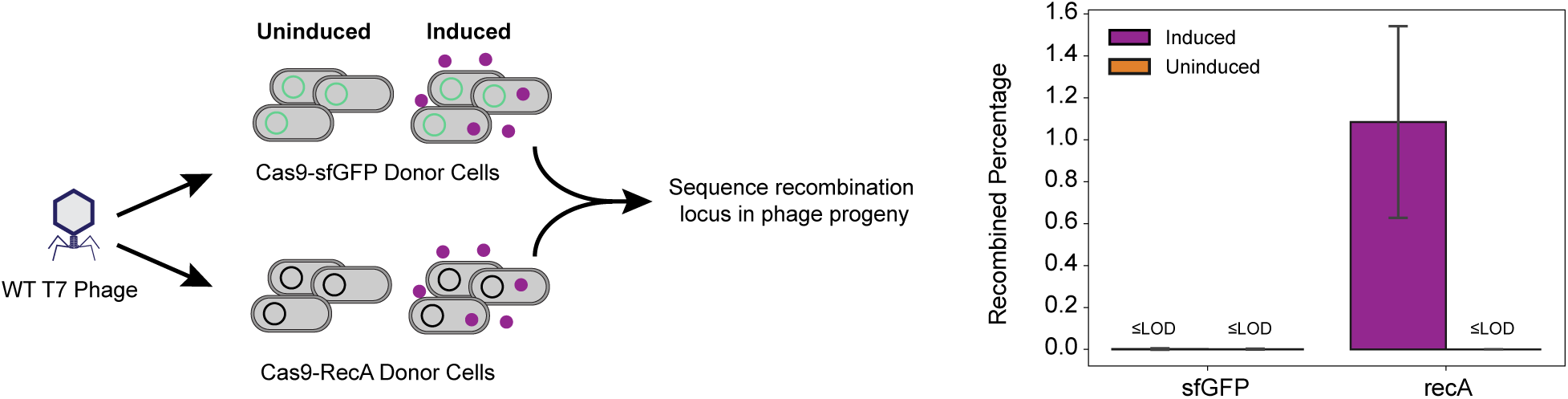
Workflow for quantifying recombination efficiency with and without recA. Cells containing donor plasmids that would replace a 100bp fragment of g1.1 in T7 with a 100bp primer binding sequence-flanked barcode were co-transformed with a Cas9-sfGFP or Cas9-RecA plasmid. Cells were either uninduced or induced with 1% anhydrotetracycline. The cultures were infected by WT T7 phage and the resulting progeny phages were used as template for PCR amplification of the target locus for sequencing. Recombined percentages were calculated as percent of total reads that contain a consensus region found in the barcode insertion. Bars represent the average (n=3 biological replicates) recombined percentages. Error bars denote standard deviation. LOD is the limit of detection which is 1 read.

**Supplementary Figure 2.**
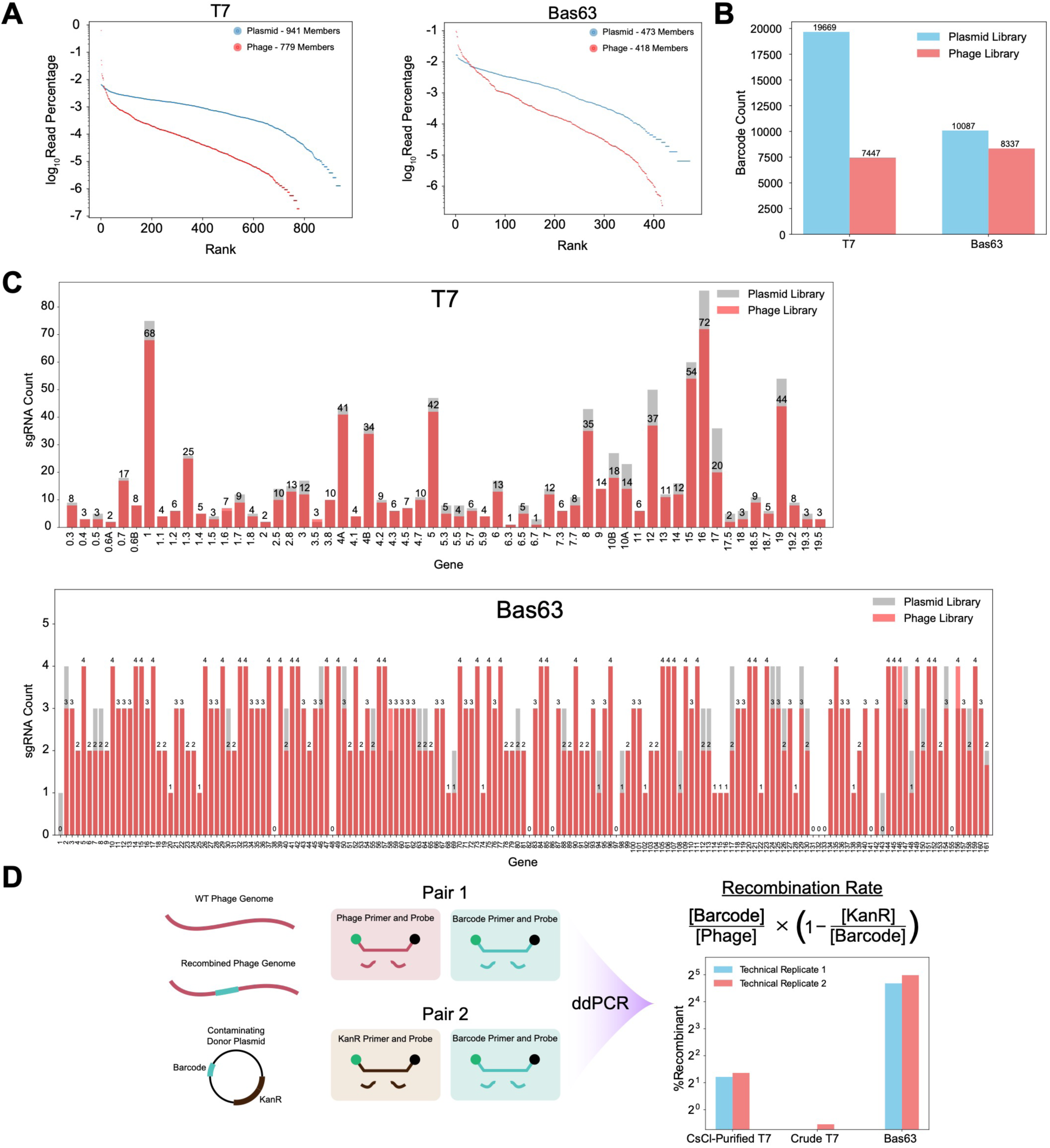
Analysis of PhageMaP library quality for T7 and Bas63. **(A)** Rank-order curves for the T7 and Bas63 library members at the donor plasmid and phage library stages. **(B)** Total barcode counts for the T7 and Bas63 library members at the donor plasmid and phage library stages. **(C)** sgRNA representation for each gene of T7 and Bas63 at the donor plasmid and phage library stages. **(D)** Estimation of recombination rate of phage libraries with digital droplet PCR (ddPCR). Low %Recombinant for “Crude T7” samples indicate high contamination of donor plasmid DNA which is largely removed after cesium chloride (CsCl) ultracentrifugation.

**Supplementary Figure 3.**
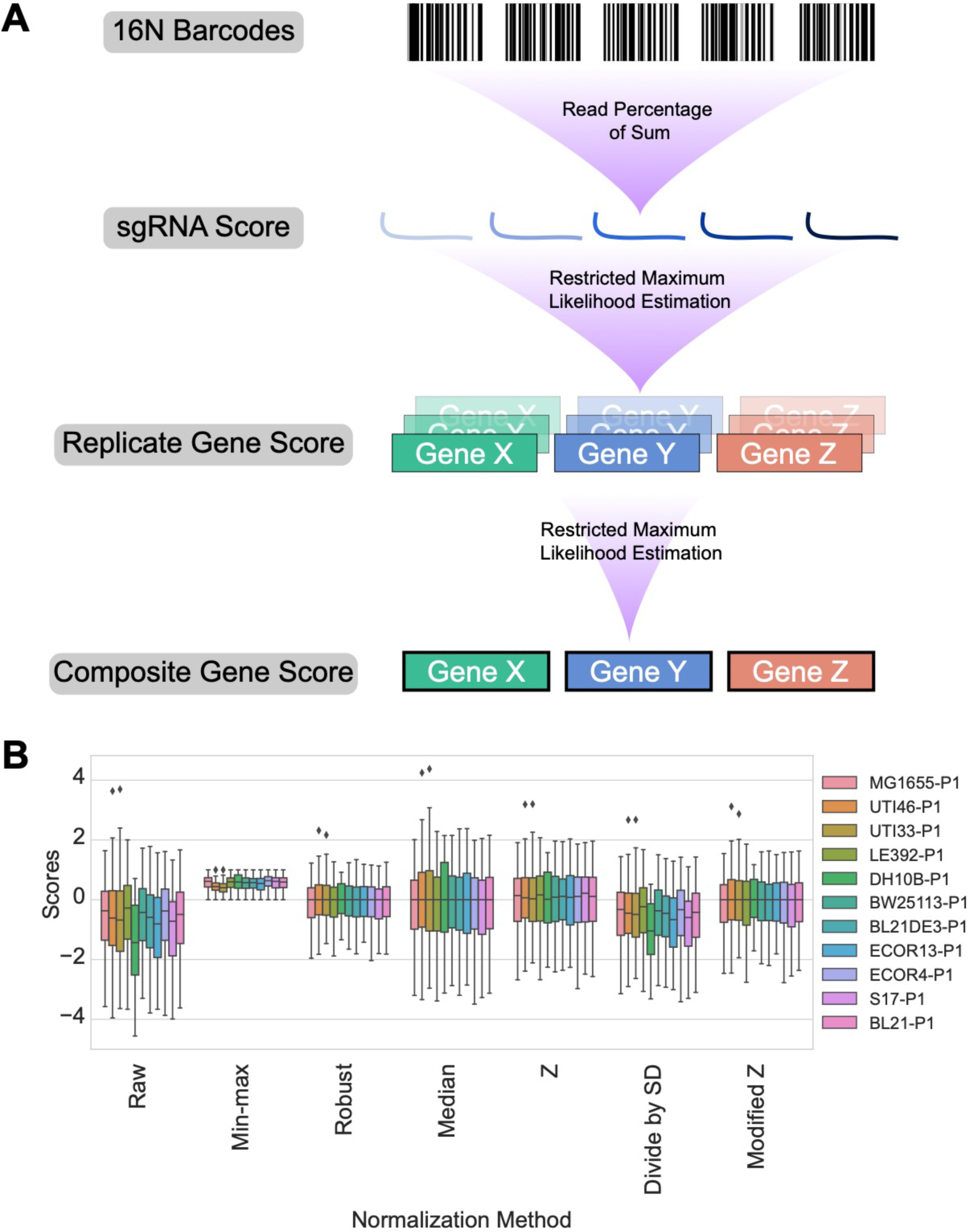
PhageMaP scoring and analysis. **(A)** Pipeline for calculating raw fitness scores from barcode counts. **(B)** Analysis of different methods for normalization. Boxplots represent the distribution of fitness scores after indicated normalization method.

**Supplementary Figure 4.**
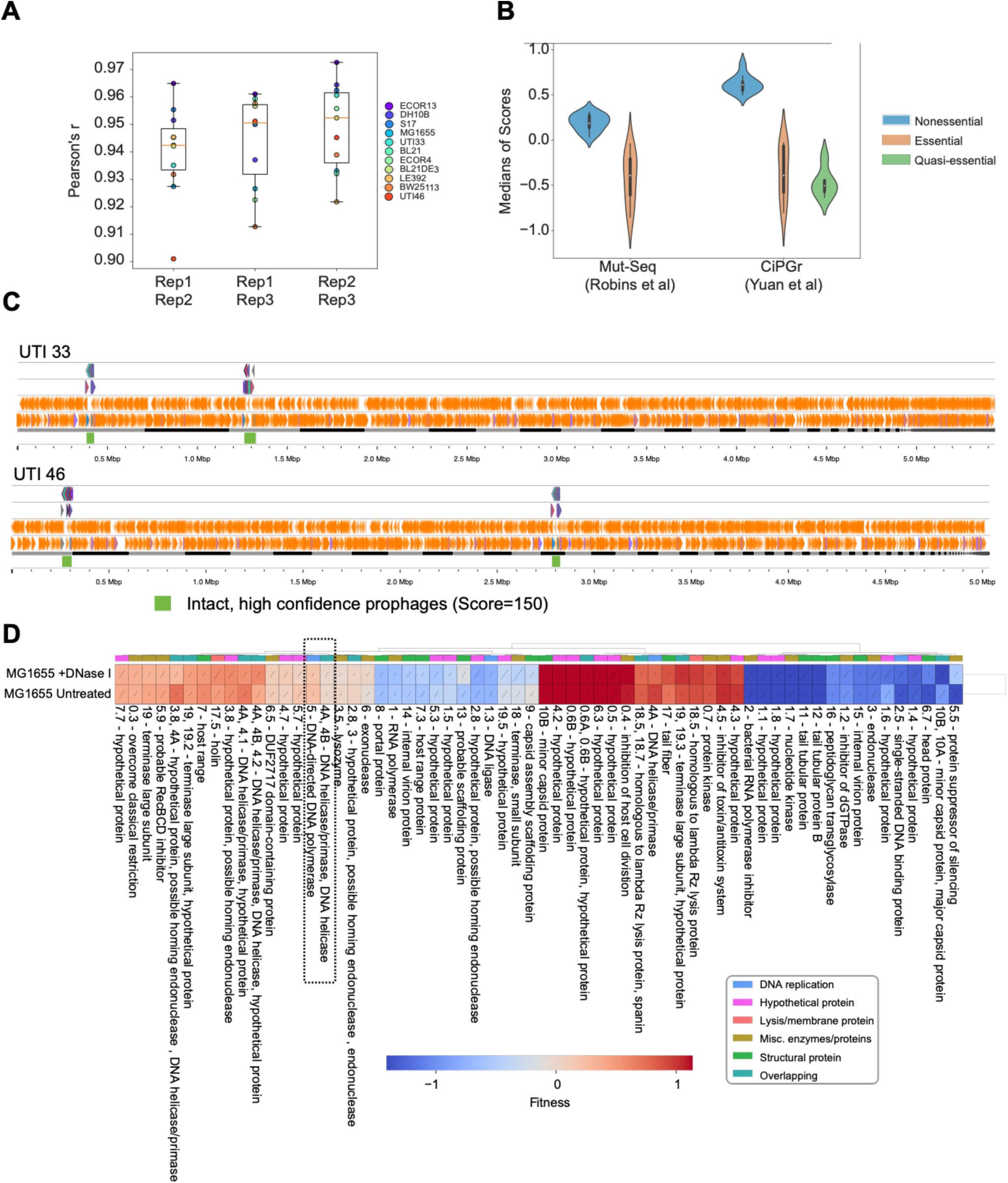
Quality of T7 PhageMaP library screening on common labaoratory *E. coli* panel. **(A)** Pair-wise Pearson’s correlation between fitness scores of replicates in each of the 11 hosts. **(B)** Comparison of PhageMaP screen results with results from previous studies with T7. The classification of each gene from the indicated study was used to group genes. The median PhageMaP score for each group was then calculated for each of the 11 hosts. The violin plot represents the median values from those 11 hosts. **(C)** Genome tracks of UTI33 and UTI46 highlighting regions with intact, high confidence prophages (green). **(D)** Heatmap of T7 fitness scores after Z-normalization for DNase-treated or untreated MG1655 lysates. A 10^th^ percentile floor and 90^th^ percentile ceiling were imposed for the color scaling. The lines through each cell represent the standard error (SE) where a full line represents SE≥1. Clustering was performed using the Euclidean metric and Ward linkage method. Metadata describes general classification of each gene. Dotted box indicates genes which were hypothesized to have different scores after DNase treatment.

**Supplementary Figure 5.**
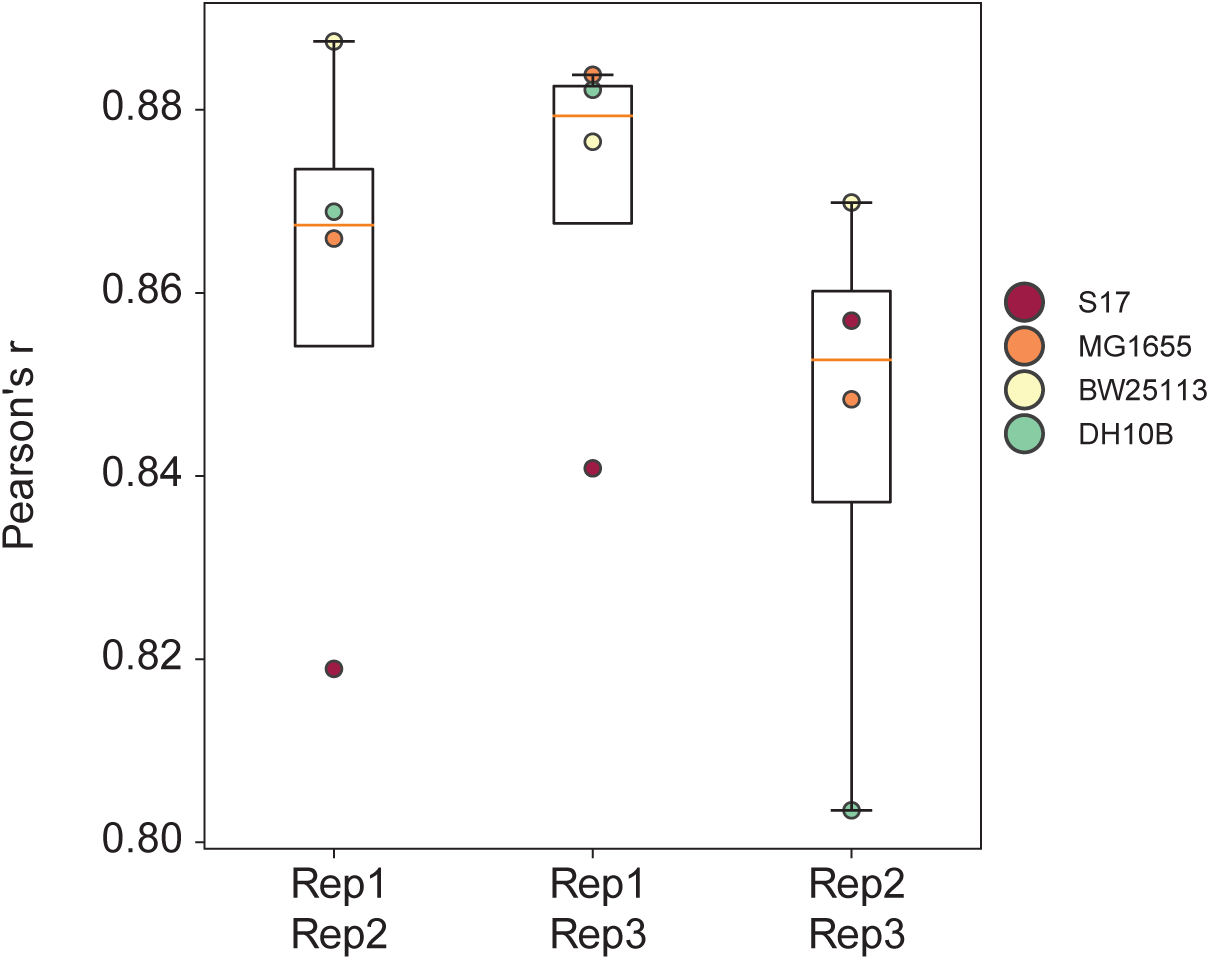
Bas63 PhageMaP reproducibility. Pair-wise Pearson’s correlation between fitness scores of replicates in the common laboratory *E. coli* panel.

**Supplementary Figure 6.**
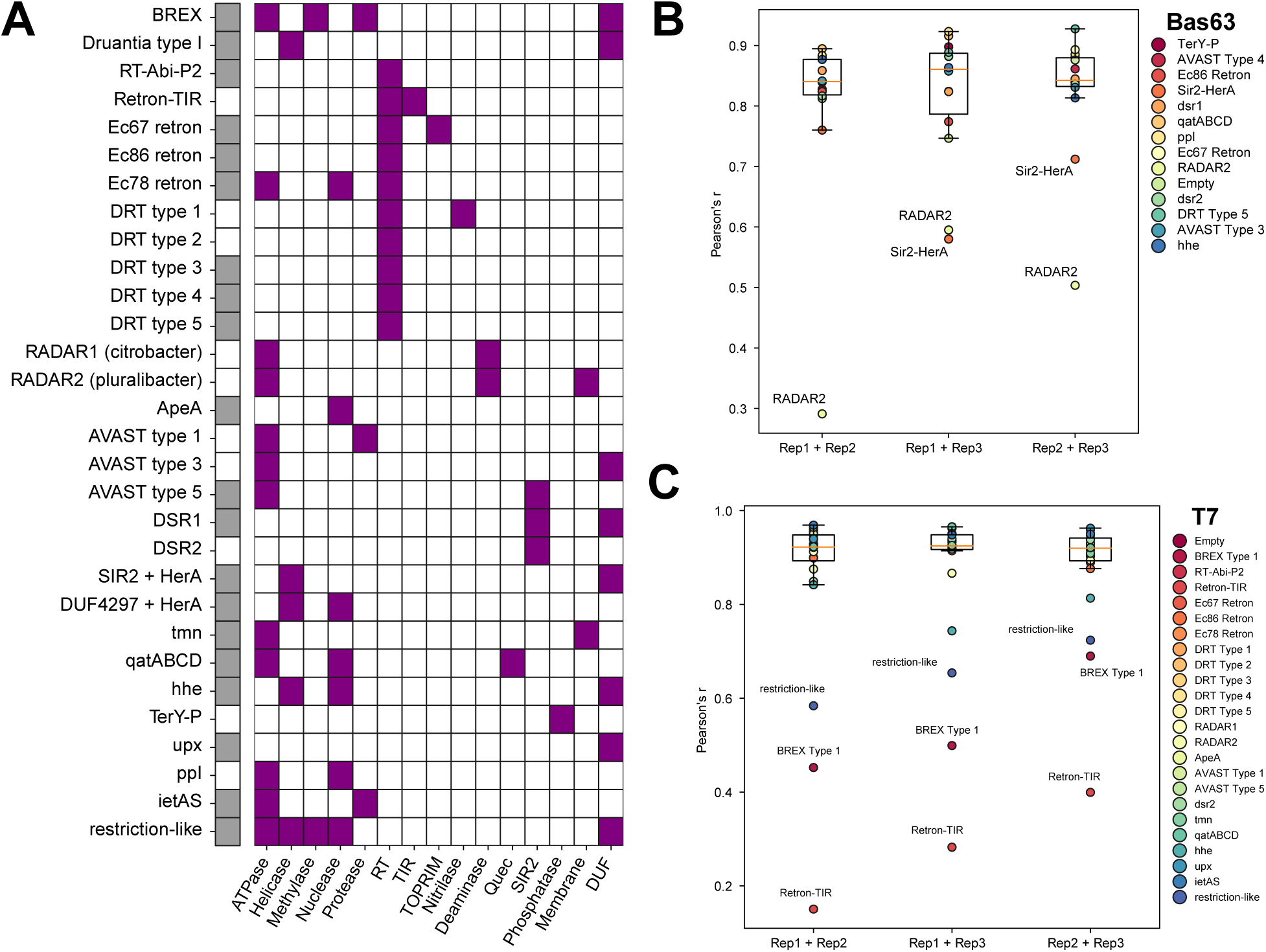
PhageMaP screen using an anti-phage defense panel. **(A)** Enzymatic domains of each defense system in tested panel. Filled colored box indicate presence of enzymatic domain. Filled gray boxes on the left indicate system is of *E. coli* origin. **(B)** Pair-wise Pearson’s correlation between fitness scores of replicates in the Bas63 screen. **(C)** Same as (B) but for T7.

**Supplementary Figure 7.**
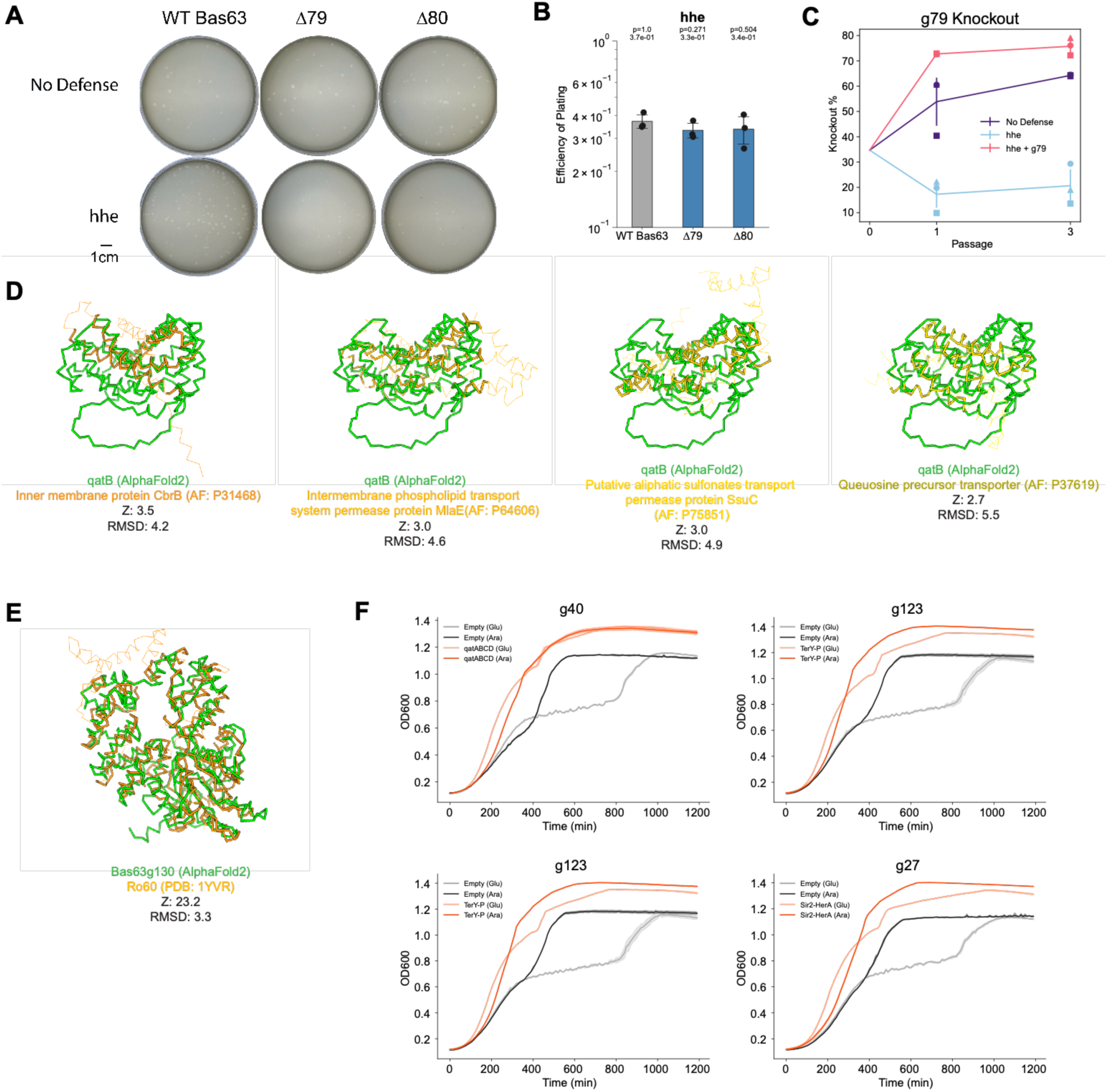
Additional analysis for Bas63 anti-phage defense screen. **(A)** Plaque morphologies of knockouts grown on the No-defense and *hhe* strains. **(B)** Efficiency of plating (EOP) measurements for Bas63 knockouts with the *hhe* host. Average titers on the control strain were first determined (3 technical replicates). Each point on the plot represents a technical replicate for EOP using those calculated averages as the denominator for each titer measurement on the defense host. Bars represent the mean of three replicates, and error bars denote the standard deviation. Hypothesis testing was performed pairwise with each knockout and WT. P-values (Welch’s t-test) and means are annotated above the bars. **(C)** Competition assay between barcoded WT Bas63 and Bas63Δ*g79* phages. Barcoded phages were mixed and passaged on the indicated hosts. Barcodes were sequenced and quantified to determine the ratio of Bas63Δ*g79* phages after each passage. **(D)** DALI structural comparisons of qatB with four membrane associated proteins. QatB (green) structure was predicted using AlphaFold2 and compared to structures from the AlphaFold Database v2 (yellow/orange). Thick yellow/orange ribbons represent aligned regions of subjects. **(E)** DALI structural comparison of Bas63gp130 (green) with Ro60 (orange). Thick orange ribbons represent aligned region of Ro60. **(F)** Additional growth curves for identified Bas63 phage defense triggers. Growth curves for validation of Bas63 phage defense triggers for different systems. Cells containing either an empty vector or the indicated defense systems were transformed with an arabinose-inducible expression construct for the specified gene. Optical density measurements were acquired every 10 minutes for 1200 total minutes. Central line represents the average of 3 replicates and the shaded flanking regions represent the standard deviation.

**Supplementary Figure 8.**
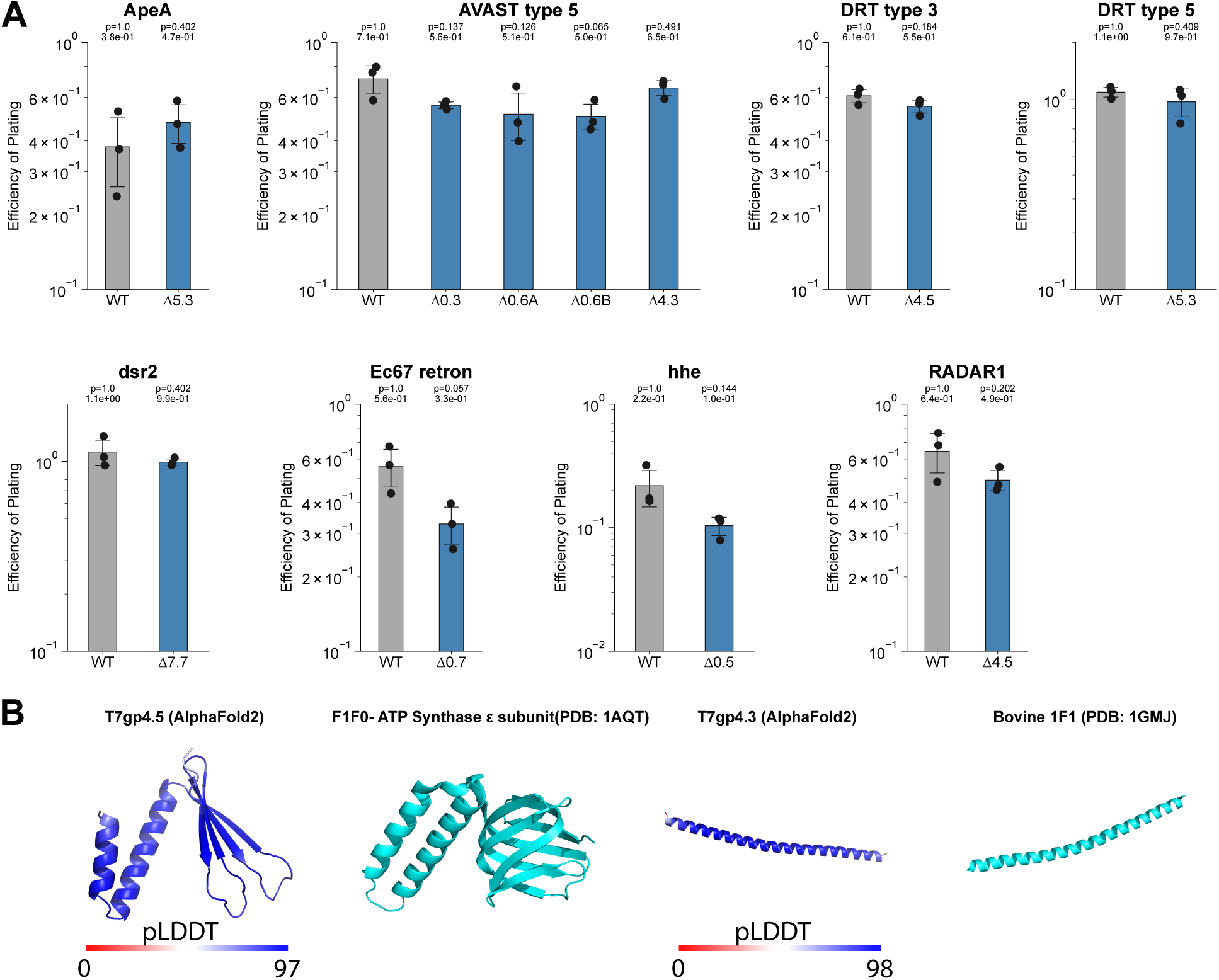
Additional analysis for counters identified from T7 anti-phage defense screen. **(A)** Additional efficiency of plating (EOP) measurements for T7 knockouts under different phage defense systems. EOP measurements for gene knockouts (potential counters) in the context of indicated phage defense system. Average titers on the empty vector were first determined (3 technical replicates). Each point on the plot represents a technical replicate for EOP using those calculated averages as the denominator for each titer measurement on the defense host. Error bars represent standard deviation. Hypothesis testing was performed pairwise with each knockout and WT. P-values (Welch’s t-test) and means are annotated above the bars. **(B)** Structural comparisons of T7gp4.5 and T7gp4.3 with the ε subunit of ATP Synthase and 1F1. T7gp4.5 and T7gp4.3 structures were predicted using AlphaFold2 and colored according to pLDDT confidence scoring. ε subunit of ATP Synthase (PDB: 1AQT) and bovine 1F1 (PDB: 1GMJ) structures were acquired from Protein Data Bank.

**Supplementary Figure 9.**
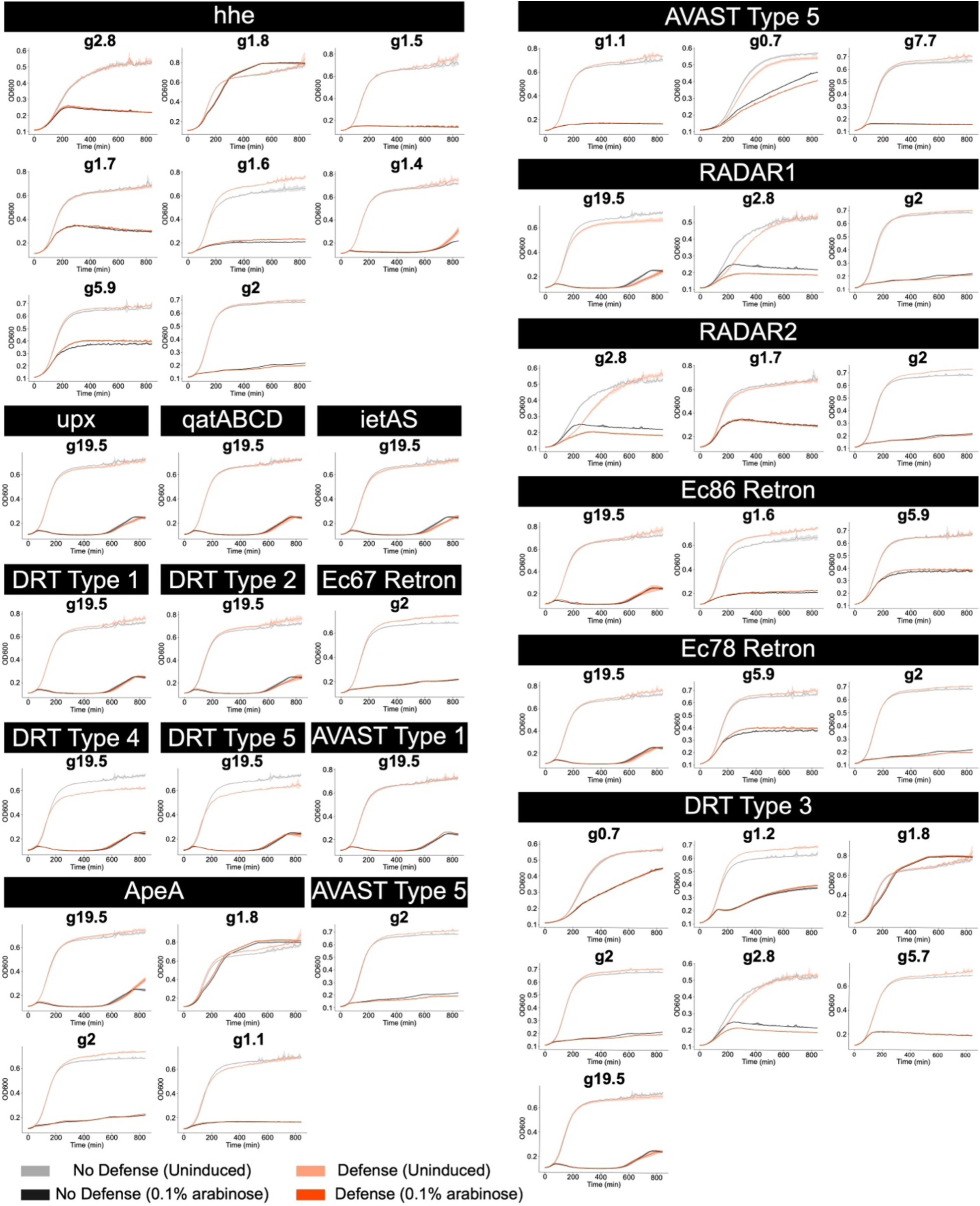
Additional growth curves for expression of T7 phage genes under different phage defense systems. Growth curves for validation of T7 phage defense triggers for different systems. Cells containing either an empty vector or the indicated defense systems were transformed with an arabinose-inducible expression construct for the specified gene. Optical density measurements were acquired every 10 minutes for 850 total minutes. Central line represents the average of 3 replicates and the shaded flanking regions represent the standard deviation.

